# In Silico Evaluation and Therapeutic Targeting of LVDD9B Protein for WSSV Inhibition: Molecular and Ecological Insights for Aquaculture Solutions

**DOI:** 10.1101/2025.04.05.647344

**Authors:** Md. Iftehimul, Neaz A. Hasan, Mst. Farzana Akter, Md. Arju Hossain, Sajia Afrin Tima, Amirul Kabir, Prottay Choudhury, Apurbo Bhowmick, Sakib Anzum Pranto, Ali Mohamod Wasaf Hasan, Siddique Akber Ansari, Md Habibur Rahman

## Abstract

**Background:** This study aimed to investigate structural dynamics, binding interactions, stability, pharmacokinetics, ecological risks, and bioactivity of shrimp receptor protein LVDD9B to identify potential therapeutic candidates against White Spot Syndrome Virus (WSSV).

**Methods:** LVDD9B protein’s 3D structure was predicted using SWISS-MODEL and validated with ProSA and Ramachandran plots. Protein-protein docking between LVDD9B and VP26 (WSSV protein) was performed using HADDOCK 2.4 server. Molecular docking, dynamics simulations, binding-free energy calculations, principal component analysis (PCA), electrostatic, and vibrational frequency analyses evaluated binding affinity, stability and polarity of complexes.

**Results:** 128-amino-acids of LVDD9B protein was predicted as predominantly cytoplasmic with stable, and hydrophilic, with structural analysis identified key secondary structures and conserved chitin-binding site. Docking studies revealed strong interactions between LVDD9B and VP26, supported by hydrogen-bonds and salt bridges. Molecular dynamics simulations demonstrated stable complexes with fluctuating RMSD values, and MM/GBSA calculations indicated favorable binding free energies. Pharmacokinetic analysis highlighted promising bioavailability and drug-like properties for Luteolin and Quercetin from *Cuscuta reflexa*, while ecological assessment identified Cosmosiin as least hazardous, with Quercetin and Luteolin showing higher toxicity. PCA revealed stable protein-ligand complexes with flexibility in Apo form. Isorhoifolin exhibited the lowest internal energy (-2099.4722 Hartree) and highest dipole moment (8.1833 Debye). Frontier orbital analysis showed HOMO-LUMO gaps (4.05–4.34 eV) influencing reactivity, while MEP and vibrational frequency analyses supported compound stability and bioactivity.

**Conclusions:** This study explores LVDD9B’s structural and interaction dynamics for developing antiviral therapy against WSSV, highlighting therapeutic potential of Cosmosiin, Isorhoifolin, Quercetin and Luteolin based on their pharmacokinetic and ecological profiles.

## Introduction

Rapid expansion of global shrimp industry, driven by increasing global demand, has led to intensive farming practices following higher stocking densities and more frequent production cycles. These practices have increased stress on shrimp populations, weakened immune responses, and created conditions conducive to the emergence and spread of disease [1], including those caused by pathogens such as viruses, bacteria, protozoa, and fungi [2, 3]. While improved culture technique, sanitation, and chemical treatments have mitigated bacteria, protozoal, and fungal diseases [4], viral diseases particularly White Spot Disease (WSD), remain a critical challenge [1, 3], causing up to 60% losses in affected population compared to 20% losses from bacterial pathogens [5]. WSD, caused by White Spot Syndrome Virus (WSSV), is a highly virulent disease responsible for catastrophic mortality in shrimp farming, with an annual economic loss estimated at US$1 billion globally with around 15% drop on world shrimp production [6]. Wide geographical distribution (China, India, East and Southeast Asia, Middle East, America, and Europe) and host range (*Penaeus monodon*, *Litopenaeus vannamei*, *Fenneropenaeus indicus*, and *Marsupenaeus japonicus*) of WSSV from both wild and farm-based source, exacerbate its impact on shrimp industry [7]. Despite the first detection of WSD in 1992 [6], efforts to develop effective antiviral therapies against this viral disease of shrimp have been hindered by limited understanding of the molecular mechanisms (specific steps, host and pathogen protein interactions) involved in propagating of WSD [8]. WSSV primarily infects shrimp via oral ingestion of infected tissues [9], crossing the basal membranes of digestive tract to reach target organs. Several proteins, including glucose transporter 1 (Glut1), C-type lectin (CL), chitin binding proteins (CBPs), peritrophin-like protein (PTs), tetraspanins, and thrombospondins facilitate viral entry and replication [10]. Apart from these proteins, cuticle proteins (CPs) including LVDD9B might be directly or indirectly involved in interaction between WSSV and shrimp cells. LVDD9B, a cuticle protein with chitin-binding domain (ChBD4), expressed in the epidermis and stomach, is implicated in the initial stages of WSSV infection. LVDD9B has shown specific binding activity with VP28, a WSSV envelope protein [11], critical for virus penetration through the shrimp cuticle. In case of LVDD9B, these CPs seem to create a more favorable environment for the virus, making them key facilitators in WSSV pathogenesis [12]. Silencing LVDD9B via RNA or dsRNA interference significantly reduces viral load and shrimp mortality at early infection stages of shrimp [13], though it cannot completely block viral replication [14], suggesting a combined role of LVDD9B and other cuticle proteins in facilitating WSSV pathogenesis. Given the limited success of antiviral strategies, including experimental vaccines especially those utilizing viral proteins like VP28 and VP26 [15], such as *Bacillus subtilis*-based oral formulations [16], research has started to concentrate on natural drugs because of their eco-accommodating and non-toxic nature. Herbal medicines such as *Ophiopogon japonicus*, geniposidic acid, coumarin, and cuminaldehyde (from *Cuminum cyminum*) have demonstrated potent antiviral properties by enhancing innate immune response and reducing oxidative stress in shrimp [17]. Dietary supplements derived from *Psidium guajava*, *Argemone mexicana* and *Gardenia jasminoides*, trigger enzymes (superoxide dismutase (SOD), alkaline phosphatase (AKP), and lysozyme), transcriptionally activates innate immune genes, increase antimicrobial peptides production, and inhibit viral replication [18–20]. However, challenges in standardizing herbal treatments, due to variability in active and inactive compounds concentrations and stability in water, limit their large-scale use in shrimp farming. Standing on this crisis of an effective therapeutic, characterizing molecular interactions between WSSV proteins and shrimp cuticle proteins is crucial. The interaction between cuticle protein LVDD9B in shrimp (*Litopenaeus vannamei*) and VP26 protein of WSSV plays critical role in viral attachment and pathogenesis. Disrupting this interaction using natural flavonoids could inhibit WSSV infection. Using in-silico modeling and molecular dynamics simulations (MDS), specific flavonoids could be identified that can bind to either LVDD9B or VP26, thereby preventing the formation of LVDD9B-VP26 complex, reducing virus’s ability to infect shrimp cells. Recent advancements in computational modeling and MDS enable detailed examination of these interactions at atomic level, offering insights into the stability, conformational changes and binding affinities of protein-protein and protein-ligand complexes. This approach may provide a foundation for developing natural, flavonoid-based therapeutic against WSSV. Several studies have mentioned the interference of variety type flavonoids in viral infections. Quercetin binds to hemagglutinin protein of influenza virus and thereby inhibits it from entering host cells, while kaempferol disrupts the Hepatitis C virus replication process by interacting with NS3 protease [21]. These findings demonstrate flavonoid-mediated inhibition of multiple viral proteins and pathways, which proves multisite inhibitory potential towards shrimp WSSV. Considering the urgent requirement for emerging therapeutic option to treat WSSV, these findings have been used as foundation for further research in proving flavonoids that can disturb the complex LVDD9B-VP26 of shrimp WSSV. In this milieu the objectives of this study have been set as threefold steps: first, to model binding interface between LVDD9B and VP26 using computational modeling techniques within a mimicked cellular context. The second is to screen the potency flavonoids of natural origin which can bind into LVDD9B and potentially inhibit viral protein of WSSV. Third, to ensure the stability and efficiency of these interactions with ligand-receptor through MDS. Lastly, analyzed the thermodynamic, chemical and geometrical properties, along with vibrational data, molecular electrostatic potentials of *Cuscuta reflexa* compounds. By achieving these objectives, we attempt to establish the basic blueprint for constructing novel bio-safe flavonoid-derived antiviral agents against WSSV in hopes of offering valuable research alternative conducive via natural product chemistry to sustain shrimp aquaculture.

## Materials and Methods

### Identification of molecular interaction between virus and host protein

The respective source of the shrimp receptor protein LVDD9B is referred to by nucleotide sequence accession number MW273781 and protein ID QQZ19802.1. It has thorough empirical analysis through bioinformatics and structural characterization directed to understand the biological significance concerning its interactions. A sequence representing the LVDD9B protein made up of 128 amino acids was downloaded in FASTA format from NCBI (https://www.ncbi.nlm.nih.gov/) and was incorporated in extensive structural predictions. Then, a set of physicochemical attributes has been established using the ProtParam tool on ExPASy (https://web.expasy.org/protparam/), such as molecular weight, amino acid composition, theoretical pI, number of positively and negatively charged residues, atomic composition, total atom number, GRAVY, extinction coefficient, aliphatic index, half-life estimate, and an instability index, all of which contribute to the understanding of the protein’s hydropathicity, stability, and functional potential [22]. CelLo (http://cello.life.nctu.edu.tw/) algorithm predictions were made for the subcellular localization, which is important in deducing the cellular roles and thus the possible protein interactions and regulatory implications [23]. Fulfilling the functional homologues of the LVDD9B protein, analyzed to NCBI BLASTP (https://blast.ncbi.nlm.nih.gov/) tools, was enabled the subsequent application of multiple sequence alignment (MSA) and phylogenetic tree generation from MEGA11 to reveal evolutionary relationships among the cuticular proteins of shrimps [24]. Secondary structure analysis was performed via PSIPRED (http://bioinf.cs.ucl.ac.uk/psipred/) and JPred (https://www.compbio.dundee.ac.uk/jpred/) to reveal the folding patterns and biological activity of LVDD9B [25, 26]. The predicted 3D model was developed based on the similarity with homologous structures from the Protein Data Bank using SWISS-MODEL for homology modelling of its chitin-binding domain 4. Subsequent validation of the model was accomplished with PROCHECK, Verify 3D, and Prosa-web for structural accuracy and by Prosa-web to get the Z-score measure of quality [27–29]. The docking results included energy scores, RMSD, and solvent-accessible surface areas are depending on protein preparation and refinement, which supported by UCSF Chimera [30]. It has been shown through protein-protein docking studies by HADDOCK 2.4, which established the interaction between the chitin-binding domain of LVDD9B and viral protein VP26 (PDB-ID: 2EDM) [31]. Finally, PDBsum (https://www.ebi.ac.uk.pdbsum) analyses elucidated the binding strength, interface characteristics, and post-translational modifications of the VP26-LVDD9B complex, offering an extensive understanding of their interaction and future avenues for drug design and experimental investigations [32]. Such integrative approaches lay down foundations in functional and structural dynamics of the LVDD9B module.

### Design a therapeutic inhibitor of LVDD9B

The ligand library construction for the current study was carried out from the India Medicinal Plants, Phytochemistry and Therapeutics (IMPPAT) database (accessed on 30 June 2024) and it comprised detailed information about 1,742,138 plants and 9,596 phytochemicals [33]. The relevant SDF files and canonical SMILES of *Cuscuta reflexa* were subsequently retrieved in preparation for analysis via PubChem (https://pubchem.ncbi.nlm.nih.gov/) database. Lastly, pharmacokinetics profiling was done to assess absorption, distribution, metabolism, excretion, and toxicity (ADMET) attributes. Physicochemical parameters were assessed using the tools SwissADME (https://www.swissadme.ch/), and PKCSM (https://biosig.lab.uq.edu.au/pkcsm/), that led to the selection of non-toxic ligands for further computational research [34]. The non-mutated tridimensional structure of the VP26 envelope protein from the WSSV (PDB ID: 2EDM) was downloaded from the Protein Data Bank (https://rcsb.org/) after the purification from metal ions, co-factors and water molecules as well as unwanted residues, and optimized with hydrogen bonds using UCSF Chimera. The active residues that contribute to the binding interactions were identified by the COACH-D (https://yanglab.qd.sdu.edu.cn/-COACH-D/) algorithm. Ligands optimized with Gasteiger method were converted to mol2 files keeping neutral charges for docking. Molecular docking preparations were conducted via PyRx 0.8 and the binding pockets were validated using site-specific analyses [35]. The docking results were saved in a CSV file with a view to the top five ligands, which were taken further for molecular interaction analyses using Ligplot+ v2.2.8 and BIOVIA Discovery Studio, emphasizing to explore hydrophobic and hydrogen bonding patterns [36]. MDS using Desmond on Linux platforms were performed for a total of 200 nanoseconds for the VP26-ligand complexes to check the stability of the system. An orthorhombic periodic bounding box, a 10 Å margin, and TIP3P water model were used to maintain the system electro-neutral with uniformly distributed ions [37]. The molecular interactions were regulated by the OPLS3e force field, and thermodynamics parameters like Root Mean Square Deviation (RMSD), Root Mean Square Fluctuation (RMSF), Solvent Accessible Surface Area (SASA), Molecular Surface Area (MoLSA), Polar Surface Area (PSA), and Radius of Gyration were analyzed. The system was maintained at an isotropic temperature of 300 K at 1 atmosphere using the Nose-Hoover thermostat and monitored at 50 ps intervals, ensuring a thermal efficiency of 1.2 kcal/mol. Simulation Interaction Diagram (SID) confirmed that the system remained stable after verifying each complex in triplicate. The MM-GBSA calculation for binding free energy was done through the platform of Inner-Schrödinger to assess affinity between ligand and strain energy. The PCA of MDS trajectory data thus effectively reduced the dimensionality of the data while at the same time preserving the information about the principal collective molecular motions in a 3Nx3N covariance matrix reconstructed configurational space in Cartesian coordination space [36]. Eigenvectors from this analysis, supported by the ‘factoextra’ package in R studio, discovered collective atomic oscillation and structural dynamics [38]. Not only did this rigorous integrative workflow characterize the phytochemicals in *Cuscuta reflexa*, but it also mechanistically elucidated their therapeutic potential against VP26 as a forward step toward potential future antiviral strategies.

### Analyzing geometrical parameters of the identified phytocompounds

The 3D structures of the identified compounds from *Cuscuta reflexa* were obtained from the PubChem database after screening. Afterwards, these structures were modified and their geometry was optimized using the Gaussian v.09 software tool [39]. The optimization process was conducted with Density Functional Theory (DFT) and was used to compute the electronic transition state. The B3LYP functional was applied in conjunction with the 6-31G (d, p) basis set, which is widely regarded as one of the most effective basis sets for analyzing thermodynamic parameters of the most interacting compounds. The chemical reactivity and other electronic parameters of the highest occupied molecular orbital (HOMO) and the lowest unoccupied orbital (LUMO), known as frontier molecular orbitals (FMOs), were evaluated using DFT analysis. This assessment involved calculating the energies of these orbitals, as well as conducting vibrational frequencies analysis, and MEP analysis [40]. The values of the vibrational frequency were multiplied by a scaling factor of 0.962 in order to ensure that they were accurate and met the requirements of the values obtained from experiments while compared with conventional references [41]. The results were observed using the GaussView 6.0 software. The density of states (DOS) plot was made using the Gauss-Sum software tool [42]. The equation has been employed to examine the properties of molecules orbitals:

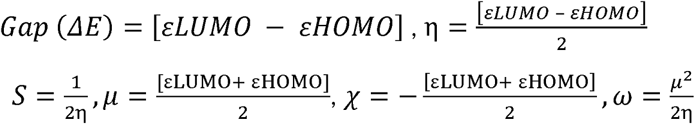

## Results

### Sequence analysis of cuticular protein

The LVDD9B protein was found to be mainly localized in the cytoplasm. Among the potential compartments, cytoplasmic localization received the highest score (2.007), strongly indicating it as the most likely site of function. Extracellular localization emerged as a secondary possibility with a score of 1.378. Predictions for other compartments, including nuclear (0.637), chloroplast (0.271), and vacuole (0.142), were substantially lower, indicating minimal likelihood. The physicochemical characteristics reveal the protein consists of 128 amino acids, with a molecular mass of approximately 14.03 kDa. It features 23 negative charged residues (Asp + Glu) and 8 positively charged residues (Arg + Lys). The protein’s aliphatic index is 66.25, points towards its decent propensity for solubility along with moderate level of thermostability, while a GRAVY score of -0.469 indicates more of its hydrophilic nature. The estimated half-life in mammalian reticulocytes in 30 hours, with longer half-lives yeast and *E. coli*. The whole sequence was divided into four separate domain such as signal peptide (1-15 Amino acids), disordered domain (48-67), Chitin binding 4 domain (30-97) and Q/N rich region (96-124). The secondary structure of the protein exhibited six distinct beta strand regions VLLDE, RYTYET, FKETV, NMVGSYRFT, VVEVRFTA and ES. Additionally, the protein contains two alpha helix domains, represented by the sequence VVLFACLIAGAF and EFVHELLAIAEQQR **(Figure S1)**. Moreover, two loops are present into the sequence such as whole-loop region (^27^DEREDQGDGN^36^) and long-loop region (^44^TPGSEGQSNMVGSYRFTDPDGNVVEVRF^67^). The top three homologs acquired from PSI-BLAST analysis of LVDD9B protein are XP_037798709.1 (*Penaeus monodon*), XP_047478664.1 (*Penaeus chinensis*), XP_042859558.1 (*Penaeus japonicus*), all showing 100% similarity to investigated protein sequence. The multiple alignment sequence (MSA) highlights a fascinating RR motif among shrimp species, with a long-loop region in the conserved sequence facilitating Ca^2+^ dependent sugars binding, compound interactions and acts a foundation for CaCO_3_ formation **(Figure 1)**.

**Figure 1:**
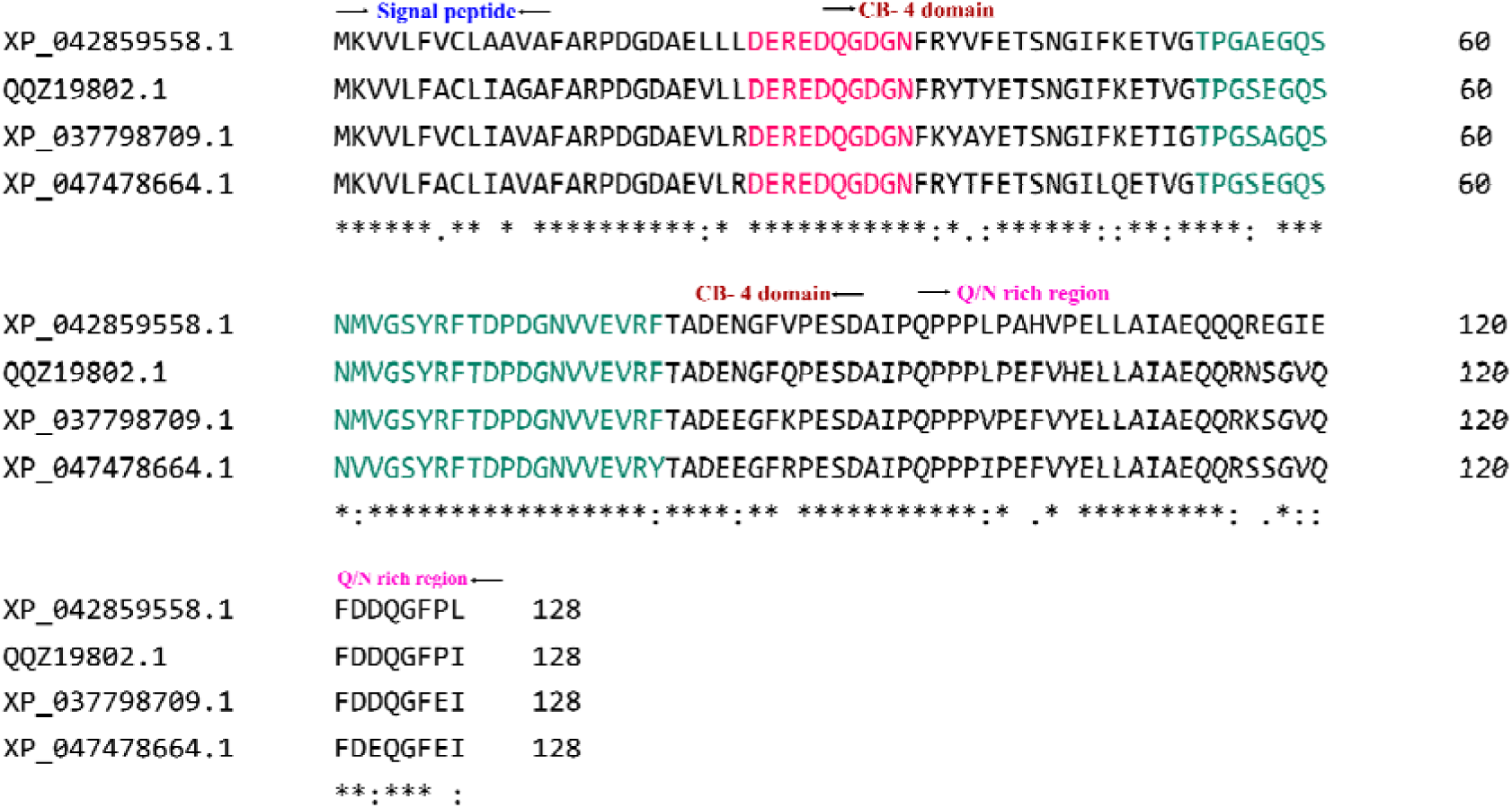
Multiple sequence alignment (MSA) of the top hits obtained from PSI-BLAST of the cuticle protein form *Litopenaeus vannamei.* The sequences are demonstrated as their protein-IDs: XP_042859558.1 (*Penaeus japonicus*), XP_037798709.1 (*Penaeus monodon*), XP_047478664.1 (*Penaeus chinensis*). Signal peptide region (1-15), Q/N rich region (96-124), Chitin binding 4 domain (30-97) are shown within arrow. The whole domain loop and long loop region is shown in magenta and turquoise color, respectively.

### Three-dimensional (3D) structure prediction and validation

The 3D structure of shrimp protein (LVDD9B) was obtained from SWISS-MODEL and depicted in (**Figure 2A**). It consists of 128 amino acids, with mostly Glycine, Glutamine and Aspartic acid present, respectively 10.2%, 9.4% and 8.6%. In addition, 26.56% of its residues form alpha helices, 19.53% form beta sheets, and 53.91% are in coil regions, paving the way for the design of novel therapeutic strategies to disrupt the infection process and protect shrimp populations from this lethal virus. The protein structural validation analysis provided a Z-score of (-2.89) in ProSa-web, indicating high quality of structure. The Verify3D plot reveals that about 53.13% of the residues exhibit a mean 3D-1D score of ≥0.1, suggesting that the model demonstrates a solid and reliable structural quality. In addition, the ERRAT demonstrates an 87.12 quality factor, indicating the model quality is good. However, the Ramachandran plot generated through the Procheck algorithm revealed that 91% of the structure’s residues were in the most favored regions, with no significant outliers. This comprehensive validation process cemented the reliability of the 3D structure (**Figure S2**); hence, the structure closely resembles the actual one and is suitable for docking with the VP26 receptor protein.

**Figure 2:**
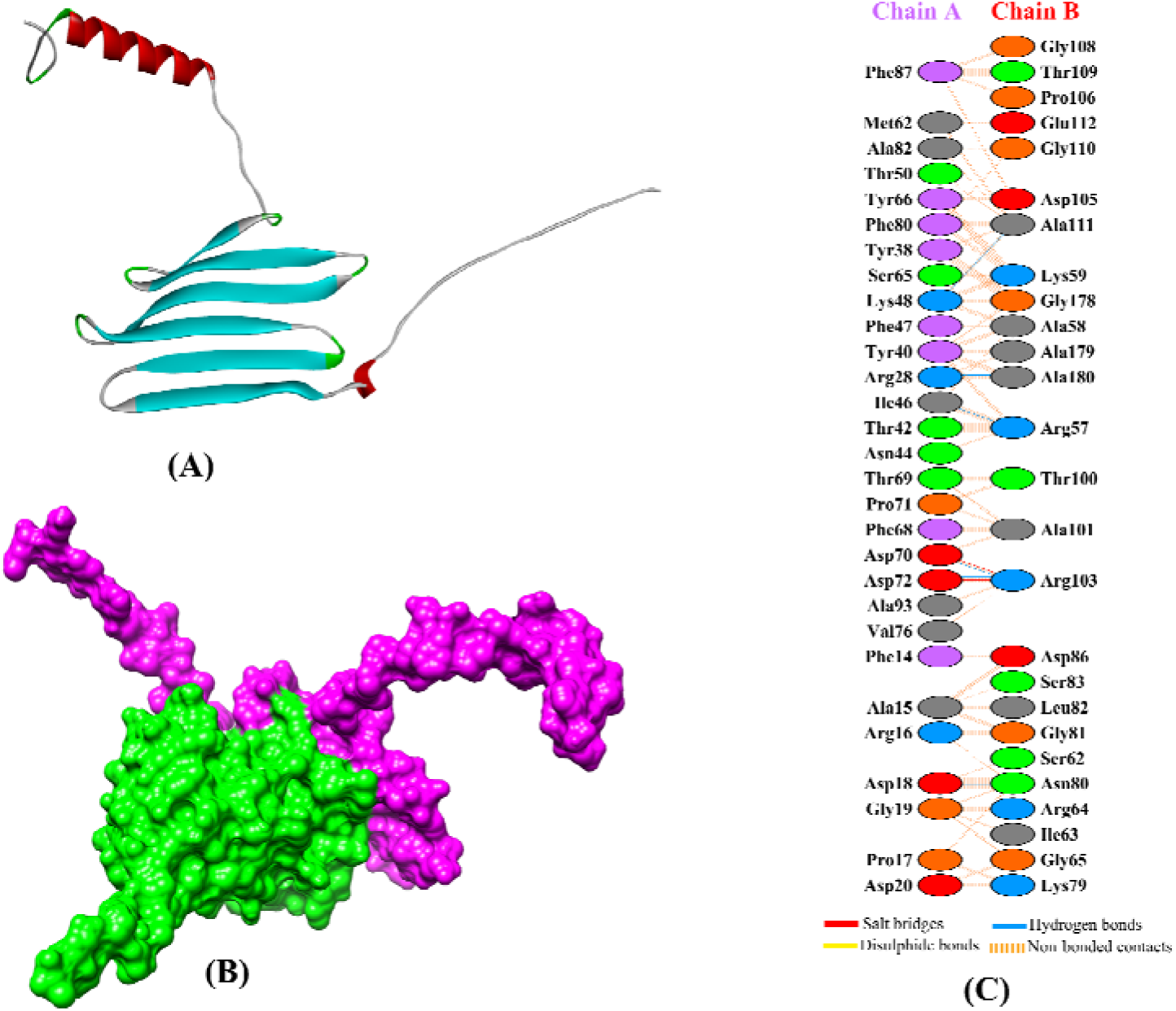
**(A)** Three-dimensional structure of the LVDD9B Protein from *Litopenaeus vannamei*, **(B)** Protein-protein docking of LVDD9B with VP26, **(C)** Protein-protein interaction between LVDD9B and VP26.

### Protein-protein docking and interpretation

The docking analysis gave us a lowest HADDOCK score of -99.9 ± 9.5 was considered as the best docked complex, indicating that the interaction between the two proteins is stable and energetically favorable. The cluster size was 18, which suggests that this particular binding mode was consistently observed in multiple simulations. The RMSD from the overall lowest-energy structure was 21.0 ± 0.1, indicating some flexibility or movement in the protein structures during the interaction, but still within an acceptable range for docking studies. In terms of energy contributions, we observed a Van der Waals energy of -79.7 ± 9.3 and an electrostatic energy of -263.1 ± 34.8, both of which are in line with what we’d expect for a stable interaction. The desolvation energy came out to -18.3 ± 3.4, which also supports the stability of the interaction in an aqueous environment. However, the restraints violation energy was relatively high at 506.9 ± 28.4, suggesting that there might be some challenges with the model, possibly due to the flexibility or particular features of the proteins involved. The Buried Surface Area (BSA) was measured at 2363.1 ± 69.7 Å², indicating a significant interaction interface. The Z-Score was -1.7, which falls within a range that usually suggests the model is of good quality when compared to the other structures in the cluster. Subsequently, we employed the PyMol molecular visualization tool to visualize our docked specimen in 3D structure. The specimen is shown in surface view mode, with the purple coloration indicating the LVDD9B protein and the green representing the VP26 protein, as depicted in **Figure 2B**.

### Protein-protein interface analysis

The interaction between LVDD9B and VP26, represented as chain A and chain B, reveals a complex interface characterized by multiple types of molecular interactions (**Figure 2C**). The analysis identifies three salt bridges, hundred and ninety-nine non bonded contacts and fourteen hydrogen bonds at the interface. Three amino acid pairs such as Arg28:Ala180, Asp72:Arg103 and Asp18:Asn80 had double hydrogen bonds between them and the rest of the pairs had single hydrogen bond existing between them, while three pairs of salt bridges were formed such as Asp20:Lys79, Asp72:Arg103, and Asp70:Arg103. These findings highlight that the stability of the protein complex is largely driven by hydrogen bonds and non-bonded contacts. The comprehensive docking and interaction studies emphasize the significant interaction between the two proteins, suggesting that this interaction could be a potential target for developing inhibitor against WSSV in shrimp.

### Determination of pharmacokinetics properties

The top lead molecules identified in this investigation, Isorhoifolin, Cosmosiin, Afzelin, Luteolin and Quercetin exhibit diverse physicochemical and pharmacokinetic characteristics. The molecular weights of these compounds range from 286.24 to 578. g/mol. Notably, Isorhoifolin has six rotatable bonds and the highest number of hydrogen bond doners (8) and acceptors (14), while Luteolin and Quercetin have the least with one rotatable bond each. These leads show topological polar surface area (TPSA) values between 111.13 to 228.97 Å², aligning with moderate drug-like parameters. The consensus Lop P values range from -0.41 to 1.73, indicating a balance between hydrophilicity and lipophilicity. Furthermore, all compounds maintain bioavailability scores of 0.55, except Isorhoifolin and more significantly violate Lipinski’s rule of five, apart from Isorhoifolin. The absorption analysis indicates that Luteolin exhibits the highest intestinal absorption (81.13%), followed by Quercetin (77.21%) and Afzelin (66.01%). Water solubility values for the phytocompounds are approximately - 3.1 to -2.5, highlighting moderate solubility, while their CaCo_2_ permeability values range from -0.229 to 0.33. In the distribution phase, Isorhoifolin has the highest volume of distribution is 1.815, while Cosmosiin shows the lowest volume 0.342. The blood-brain barrier (BBB) permeability values reveal that Luteolin has the best BBB penetration (-0.907), whereas Isorhoifolin has the poorest -1.72. CNS permeability values remain within acceptable range from -4.727 to -2.251, implying satisfactory CNS distribution. The metabolism analysis demonstrates that all compounds neither actively inhibited the metabolic enzyme nor acted as specific substrates for CYP2D6 or CYP4A4. Although compounds are usually metabolized with the help of CYP2C19, CYP2C9, CYP2D6, and CYP3A4 enzymes, most of these compounds did not inhibit any of these enzymes, while Luteolin exhibits inhibitory activities in CYP2C9. None of the compounds are renal OCT2 substrates, indicating efficient elimination pathways. Finally, all bioactive compounds demonstrated no toxicity on AMES toxicity or hepatotoxicity. Besides, the toxicity profiles for oral rat chronic, skin sensitization and tolerated dose are presented in **Table S2.**

### Ecological risk assessment

Bioactive compounds from *Cuscuta reflexa* such as Isorhoifolin, Cosmosiin, Afzelin, Luteolin, and Quercetin reveals distinctive patterns of environmental impact, particularly with regard to toxicity and biodegradability. Cosmosiin is not hazardous to a number of important species found in aquatic environments, such as *Daphnia magna* (0.10), fish (0.34), *Pseudokirchneriella subcapitata* (0.57), and crustaceans (0.12). This implies that Cosmosiin is comparatively safe for the ecological environment. Comparatively, Luteolin and Quercetin consistently rank as the most toxic compounds, particularly in *T. pyriformis* (0.98 and 0.97, respectively) and *P. subcapitata* (0.82). Furthermore, certain fish species, such as rainbow trout, bluegill sunfish, and fathead minnow, exhibit increased susceptibility to quercetin and luteolin, with reported toxicity values for rainbow trout reaching as high as 0.67. Yet again, Cosmosiin shows no toxicity for terrestrial organisms; this is especially true for honeybees (0.05), quail species like *Colinus virginanus* (0.03), and ducks (*Anas platyrhynchos*) (0.03). In contrast, all of the investigated compounds exhibited comparatively no toxicity to these species, as measured by values of 0.09, 0.10, and 0.15 for ducks, quail, and honey bees, respectively. According to these findings, compounds may be safer for pollinators and bird species alike. The bioconcentration factor (BCF) and biodegradability measures offer further information regarding environmental persistence. The bioconcentration potential of Cosmosiin is the lowest (0.008), suggesting a lower chance of bioaccumulation in living things. On the other hand, Quercetin has the greatest BCF (0.07), suggesting that it has a higher potential to accumulate in the food chain. Additionally, Quercetin has the lowest biodegradability (0.17) compared to Cosmosiin, which is moderate (0.27), indicating that Quercetin may remain in the environment longer and present longer ecological concerns. While Isorhoifolin is the most biodegradable compound (0.37), it shows intermediate toxicity in aquatic species like fish (0.53) and *D. magna* (0.20), suggesting a speedier breakdown under environmental conditions and potentially mitigating long-term ecological impact (**Table 1)**.

**Table 1:** Ecological risk assessment of the most interacting therapeutic compounds. toxicity.

| Properties | Action | Isorhoifolin | Cosmosiin | Afzelin | Luteolin | Quercetin |
| --- | --- | --- | --- | --- | --- | --- |
| Aquatic organism | *P. subcapitata* toxicity | 0.33 | 0.57 | 0.80 | 0.82 | 0.82 |
|  | Crustaceans' toxicity | 0.25 | 0.12 | 0.27 | 0.42 | 0.46 |
|  | *D. magna* toxicity | 0.20 | 0.10 | 0.26 | 0.41 | 0.46 |
|  | Fish toxicity | 0.53 | 0.34 | 0.56 | 0.57 | 0.53 |
|  | Fathead minnow toxicity | 0.25 | 0.23 | 0.38 | 0.53 | 0.51 |
|  | Bluegill sunfish toxicity | 0.32 | 0.21 | 0.46 | 0.65 | 0.57 |
|  | Rainbow trout toxicity | 0.37 | 0.24 | 0.51 | 0.67 | 0.67 |
|  | Sheepshead minnow toxicity | 0.27 | 0.23 | 0.36 | 0.62 | 0.60 |
|  | *T. pyriformis* toxicity | 0.40 | 0.90 | 0.87 | 0.98 | 0.97 |
| Terrestrial organism | Honey bee toxicity | 0.22 | 0.05 | 0.06 | 0.07 | 0.09 |
|  | *Colinus virginianus* toxicity | 0.08 | 0.03 | 0.05 | 0.08 | 0.10 |
|  | *Anas platyrhynchos* toxicity | 0.09 | 0.03 | 0.06 | 0.12 | 0.15 |
| Other | Bioconcentration Factor | 0.02 | 0.008 | 0.02 | 0.05 | 0.07 |
|  | Biodegradability | 0.37 | 0.27 | 0.25 | 0.30 | 0.17 |

### Site-specific docking and post docking interactions

Binding energy indicates the strength and stability between compound and the target protein. Among our tested compounds Isorhoifolin, Cosmosiin, Afzelin, Luteolin, and Quercetin shows varying docking score against the target protein (LVDD9B). The binding energies for Isorhoifolin, Cosmosiin, Afzelin, Luteolin, and Quercetin were found to be -8.5 Kcal/mol, -8 Kcal/mol, -7.3 Kcal/mol, -7.2 Kcal/mol and 7.1Kcal/mol, respectively. The more negative the binding energy, the stronger the interaction between the ligand and the target protein. The analysis of non – covalent interactions (hydrogen and hydrophobic bonds) was conducted using the Ligplot+ (V 2.2.8) program. Isorhoifolin and the LVDD9B protein exhibit a strong binding interaction. The residues Phe14 (3.00 Å), Pro71 (3.21 Å), Tyr40 (2.85 Å), and Asn44 (2.84 Å and 2.87 Å) were the five hydrogen bonds it formed. In addition, the residues Ala13, Asp18, Asp70, Asp72, Gly73, Ile46, Phe68, Thr69, and Tyr66 form hydrophobic interaction with Isorhoifolin. Additionally, Asp70 (2.93 Å), Pro71 (3.20 Å), Tyr40 (2.76 Å), and Asn44 (3.01 Å and 2.92 Å) established five hydrogen bonds with Cosmosiin and Amino acid residues Tyr66, Phe68, Thr69, Asp18, and Ile46 are involved in the hydrophobic interactions. Afzelin interacts with the LVDD9B proteins amino acid residues and forming five hydrogen bonds with the following residues: Asp70 (3.14 Å), Pro17 (3.24 Å), Asp20 (3.25 Å), Asn44 (2.86 Å), and Thr69 (3.01 Å). Additionally, hydrophobic interactions were observed with Arg16, Asp18, Gly45, Gly73, Pro71, and Ser43. Furthermore, Luteolin_LVDD9B complex exhibited 5 hydrophobic bond (Pro71, Arg16, Asn44, Ser43, Asp18) and fitted with 5 hydrogens bonds with the LVDD9B protein involving the amino acid residues such as Asp20, Pro17, Thr69, Asp70. Moreover, quercetin demonstrated hydrogen bonding interactions with the LVDD9B protein’s amino acid residues at Thr69 (2.93 Å), Asp20 (3.19 Å), Pro17 (3.14 Å), and Asp70 (3.07 Å). The receptor of the LVDD9B protein established hydrophobic interaction with quercetin involving Pro71, Asp18, Arg16, Ser43, Asn44 amino acid residues of the protein **(Table S1** and **Figure 3)**.

**Figure 3:**
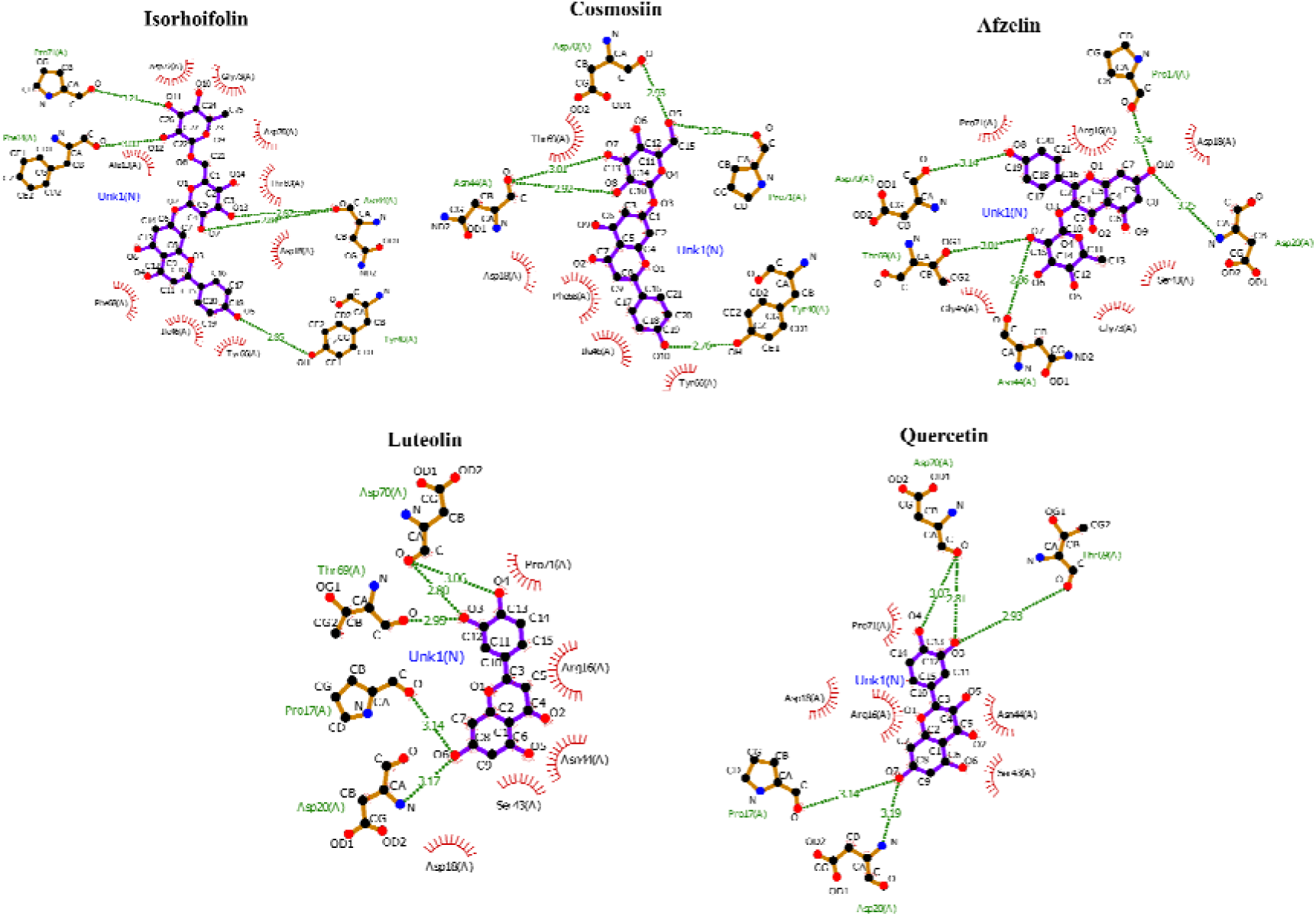
Graphical 2D representation of post docking interactions between the target protein LVDD9B and phytochemicals derived from *Cuscuta reflexa*.

### Molecular dynamics simulation study

The stability and intermolecular interactions of the system are often evaluated in a real-time environment in MDS of protein-ligand complexes. From the output trajectories, we analyzed six critical parameters such as root mean square deviation (RMSD), root mean square fluctuation (RMSF), radius of gyration (rGyr), solvent-accessible surface area (SASA), molecular surface area (MolSA), and polar surface area (PSA). In RMSD analysis, our tested ligand molecules show changes when interacting with the LVDD9B hypothetical protein. Quercetin and Luteolin exhibit the highest RMSD values of 23.96 Å and 21.31 Å, whereas Isorhoifolin, Cosmosiin and Afzelin shows 19.75 Å, 19.81 Å and 20.60 Å, respectively (**Figure 4A**). The RMSD values of the phytocompounds fluctuated beyond the acceptable range due to the hypothetical nature of the protein structure, whereas the ligand RMSD values stayed within the acceptable range. Furthermore, the RMSF value indicates the changes of protein chains during their interaction with ligand molecules. There were no significant changes observed in the RMSF curves for investigated compounds. However, while the other compounds showed average values of approximately 16 Å with a chance of fluctuating during the simulated environment, Luteolin exhibits a notable value 10.82 Å **(Figure 4B)**. A larger radius of gyration (Rg) implies that the structure is more spread out and dynamic, whereas a smaller Rg signifies a more condensed and tightly packed structure. In our results, Rg values for Isorhoifolin, Cosmosiin, Afzelin, Luteolin, and Quercetin were showed 6.0, 5.26, 4.32, 3.93 and 3.84 Å, respectively (**Figure 4C**). This is one of the most crucial factors in a biomolecule’s structural function prediction due to its ability to demonstrate fluctuations in intricate density over time. The solvent accessible surface area (SASA) significantly impacts the stability of protein and biomolecules. Throughout this research, the SASA values for the selected ligands range from 2 to 500 Å, indicating that the amino acid residues of the protein structure were predominantly exposed to the chosen ligand molecules (**Figure 4D**). Molecular surface area, or simply MolSA, refers to the equivalent of van der Waals surface area estimated with a 1.4 probe radius. In this in-silico study, all investigated phytochemicals displayed van der Waals behavior; the recorded values are 491.24 for Isorhoifolin, 374.90 for Cosmosiin, 363.33 for Afzelin, 254.08 for Luteolin and 260.26 for the Quercetin (**Figure 4E**). Lastly, in Polar surface area (PSA) analysis, Isorhoifolin presents highest surface area approximately 422 Å (**Figure 4F**). In the simulation, intermolecular interactions formed multiple bonds, such as hydrogen bonds, hydrophobic bonds, water bridge connections and so on (**Figure S3**), all of which are important for stabilizing the compound with the receptor protein. The MDS analysis revealed that most of the examined compounds showed strong hydrogen bonds and hydrophobic bonds were also prominent during the simulation.

**Figure 4:**
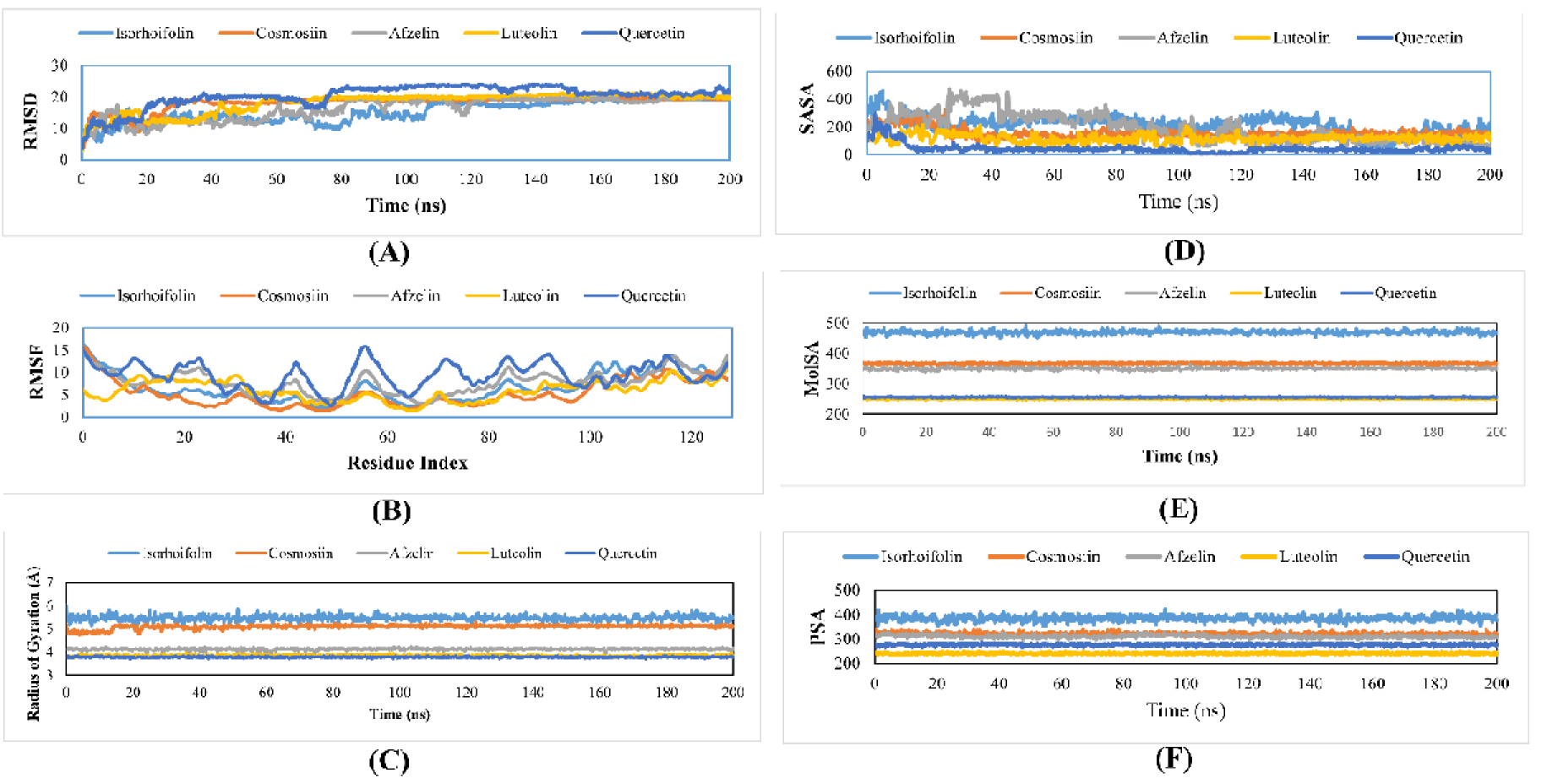
The (A) Root Mean Square Deviation (RMSD, B) Root-mean Square Fluctuation (RMSF). (C) Radius of Gyration, (D) solvent accessible surface area (SASA) (E) molecular surface area (MolSA) and (F) polar surface area (PSA) of the protein-ligand complex compounds analyzed 200ns simulated interaction diagrams.

### MM-GBSA calculation

The analysis of seven parameters showed that all compounds exhibited two positive and five negative binding free energy values. Among them, Luteolin emerged as the compound with the most positive values in several parameters. The dG_Bind_Covalent value was positive for all compounds, indicating that covalent bonds formation doesn’t occur spontaneously. Likewise, the positive dG_Bind_Solv_GB indicated that none of the compounds would dissolve easily in aqueous environments, as they require more than 20 kcal/mol. Among the five negative parameters, dG_Bind_Hbond displayed the least energy release, followed by dG_Bind_Lipo and dG_Bind_Coulomb, which confirmed lipophilicity and spontaneous electrostatic interactions, respectively. Lastly, negative value of dG_Bind_vdw showed strong van der Waals interactions expect Luteolin, and the highly negative dG_Bind value emphasized spontaneous receptor-ligand binding (**Figure 5**).

**Figure 5:**
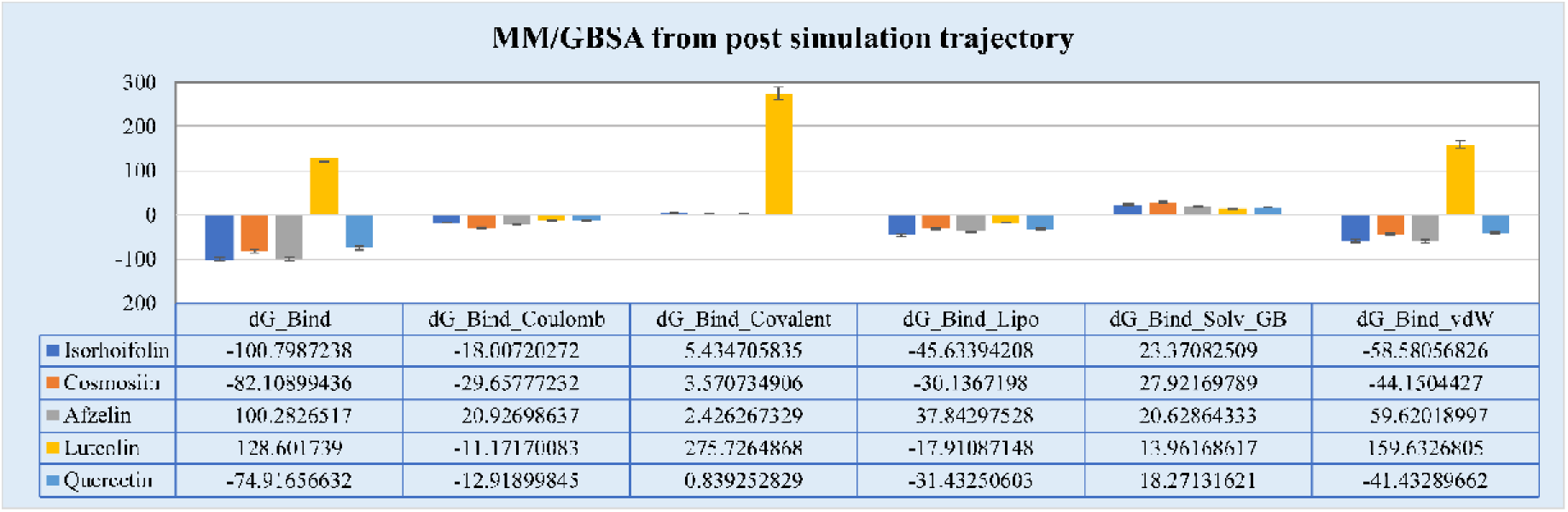
Determine the several parameters of MM-GBSA for Phytochemicals.

### Principal component analysis

In our study, Principal component analysis (PCA) was performed to explore the conformational flexibility and diversity of conformations that emerged from the stable trajectory obtained from the 200 ns MDS. The maximum collective motion is captured by the first 50 eigenvectors or principal components (PCs). It is observed that the Apo form shows diversity of conformation during the simulations (-13 to 2). However, the Luteolin–LVDD9B complex shows similar diversity of conformations during simulation (-13 to 2) and the other protein-ligand complex distribution of conformations changes during simulation (-9 to 2). In addition, the PCA plot shows that apo-protein and protein-ligand complexes motion clusters are tightly distributed (-6 to 2) and occupy less space, which indicates a more stable complex, whereas those that occupy more space indicate a less stable protein-ligand complex. In these studies, it reveals that our examined ligands with protein are well equilibrated and stabilized during the simulation (**Figure 6**).

**Figure 6:**
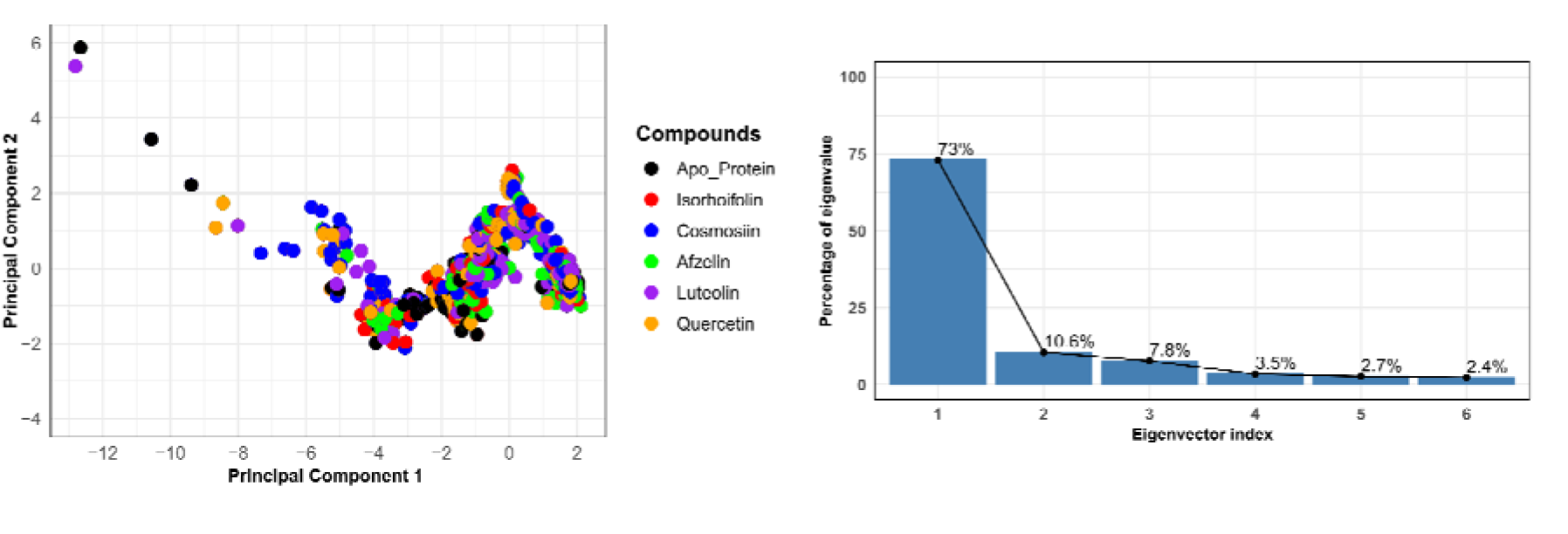
Principal component analysis of the best interacting bioactive compounds.

### Thermodynamic properties

The comprehensive analysis of the molecular and thermodynamic properties of phytoconstituents from *Cuscuta reflexa* provides significant insights into their potential biological activities and therapeutic applications. Among the compounds evaluated, Isorhoifolin, characterized by its molecular formula C[[H[[O[[ and a substantial molecular weight of 578.5 g/mol, stands out due to its remarkably low internal energy of - 2099.4722 Hartree **(Figure 7A** and **Table 2)**. This indicates a highly stable molecular structure, which may confer enhanced resilience against metabolic degradation, thereby prolonging its bioavailability and efficacy in biological systems. Additionally, Isorhoifolin’s dipole moment of 8.1833 Debye suggests a high degree of polarity, facilitating strong intermolecular interactions with polar biological targets such as proteins and enzymes, potentially enhancing its anticancer properties **(Figure 7B)**. In comparison, Cosmosiin and Afzelin, both with molecular weights of 432.4 g/mol, display similar energetic characteristics but differ in their dipole moments, which may affect their interaction dynamics. Luteolin (C[[H[[O[), despite its lower molecular weight of 286.24 g/mol, possesses an internal energy of -1028.7454 Hartree, indicating moderate stability, while its dipole moment of 4.9918 Debye points to moderate polarity. This could imply distinct mechanisms of action compared to more polar compounds. Quercetin, with a dipole moment of 3.4719 Debye, exhibits the lowest thermodynamic stability (-1104.0123 Hartree) among the compounds assessed, suggesting that its interactions may be predominantly hydrophobic in nature.

**Figure 7:**
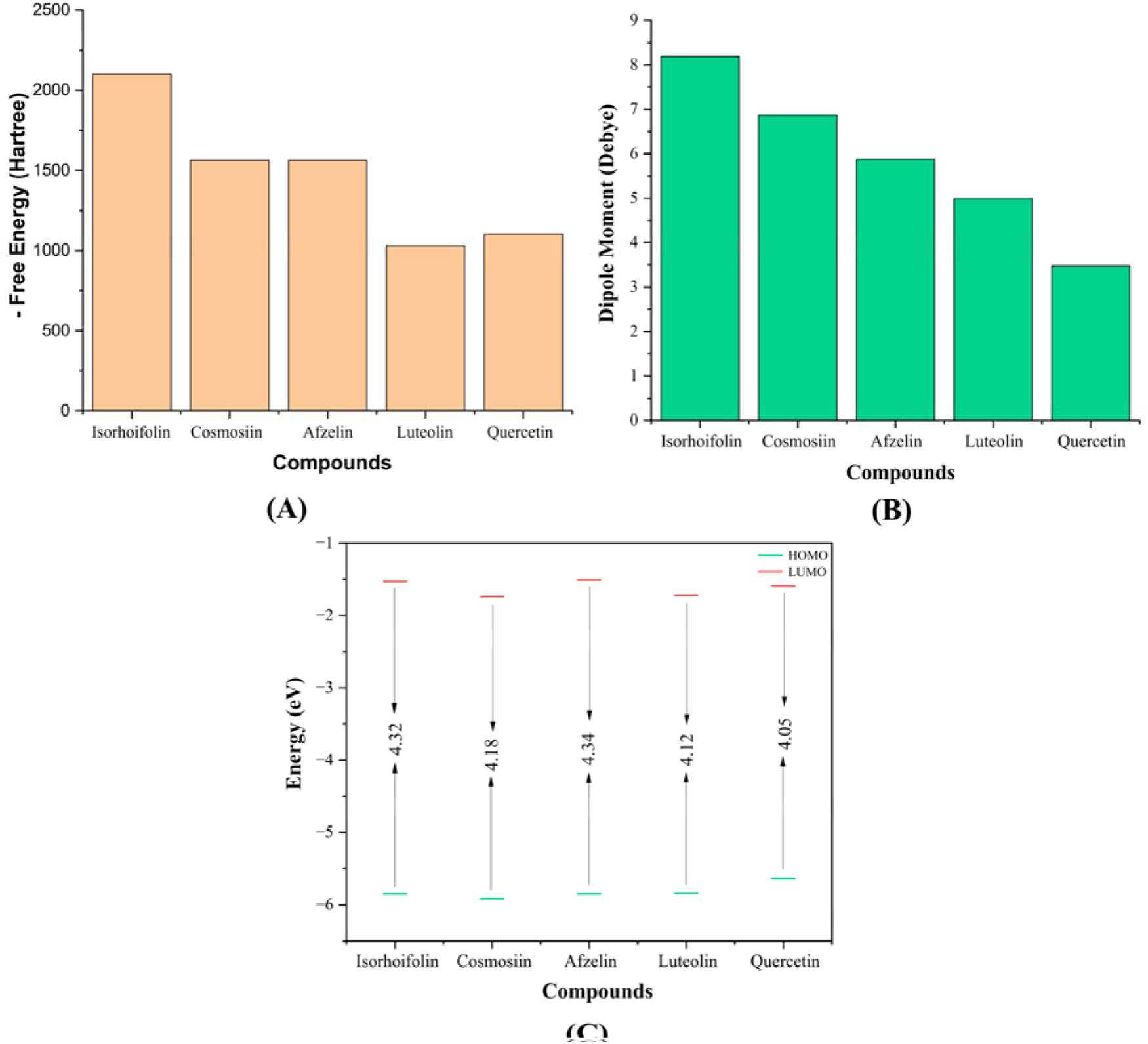
Thermodynamic properties of selected bioactive compounds from *Cuscuta reflexa* (A) Free energy, (B) Dipole moment and (C) Energy gap of HOMO-LUMO.

**Table 2:** Molecular formula (MF), molecular weight (MW), energies (Hartree), and dipole moment (Debye) of phytoconstituents of *Cuscuta reflexa*.

| Name | Molecular formula | Molecular Weight | Internal Energy (Hartree) | Enthalpy | Gibbs Free Energy | Dipole Moment (Debye) |
| --- | --- | --- | --- | --- | --- | --- |
| Isorhoifolin | C27H30O14 | 578.5 | -2099.4722 | -2099.4712 | -2099.5801 | 8.1833 |
| Cosmosiin | C21H20O10 | 432.4 | -1564.1003 | -1564.0994 | -1564.1865 | 6.8647 |
| Afzelin | C21H20O10 | 432.4 | -1564.0775 | -1564.0765 | -1564.1649 | 5.8739 |
| Luteolin | C15H10O6 | 286.24 | -1028.7454 | -1028.7445 | -1028.8069 | 4.9918 |
| Quercetin | C15H10O7 | 302.23 | -1103.9486 | -1103.9477 | -1104.0123 | 3.4719 |

### Frontier molecular orbital analysis

The HOMO-LUMO calculated energy gap of the selected phytocompounds range between 4.05 to 4.34 eV. Quercetin exhibits the smallest HOMO-LUMO gap of 4.05 eV, indicating its high reactivity and potential for electronic transitions. In contrast, Afzelin has the largest gap of 4.34eV, reflecting greater stability and lower reactivity. Compounds such as Cosmosiin, Isorhoifolin, and Luteolin show intermediate gaps such as 4.18, 4.32 and 4.12 **(Figure 7C)**, while Cosmosiin shows the highest electrophilicity (3.50 eV), suggesting strong electron-accepting properties. Additionally, Cosmosiin has the highest electronegativity (3.83 eV), further supporting its electron affinity. In contrast, Afzelin is the hardest molecule (η = 2.17 eV), implying greater resistance to electronic deformation, whereas Quercetin is the softest (S = 0.4938 eV), indicating higher chemical reactivity (**Table 3**). The efficiency of kinetics and the impact on protein interactions are heavily influenced by the HOMO and LUMO orbitals. These orbitals represent regions with high and low electron densities, respectively. According to **Figure S4**, LUMO is often situated in cation regions where positive charges are observed, resulting in an extra electron being added to the LUMO. In contrast, HOMO is generally found in anion regions, where the negative charge is present. Furthermore, the Density of state (DOS) analyzed to understanding more deeper about HOMO-LUMO and their gap, the plot depicted on **Figure 8**. The band gap of each molecule is represented by arrows in the figure, color-coded for easy comparison. Quercetiin, with the smallest energy gap of 4.05 eV, stands out as the most reactive but also the least stable molecule. The energy gaps for Isorhoifolin, Cosmosiin, Afzelin, and Luteolin were 4.32 eV, 4.18 eV, 4.34 eV and 4.12 eV, respectively, fall within the mid-range, indicating moderate reactivity and stability.

**Table 3:** Energy (eV) of HOMO-LUMO, gap, hardness (η), (S), chemical potential (μ), electronegativity (χ), and electrophilicity (ω) of *Cuscuta reflexa* compounds.

| Name | $E_{HOMO}$ | $L_{HOMO}$ | $E_{gap}$ | $\eta$ | $S$ | $\mu$ | $\chi$ | $\omega$ |
| --- | --- | --- | --- | --- | --- | --- | --- | --- |
| Isorhoifolin | -5.85 | -1.53 | 4.32 | 2.16 | 0.4629 | -3.69 | 3.69 | 3.15 |
| Cosmosiin | -5.92 | -1.74 | 4.18 | 2.09 | 0.4784 | -3.83 | 3.83 | 3.50 |
| Afzelin | -5.85 | -1.51 | 4.34 | 2.17 | 0.4608 | -3.68 | 3.68 | 3.12 |
| Luteolin | -5.84 | -1.72 | 4.12 | 2.06 | 0.4854 | -3.78 | 3.78 | 3.46 |
| Quercetin | -5.64 | -1.59 | 4.05 | 2.025 | 0.4938 | -3.61 | 3.61 | 3.22 |

**Figure 8:**
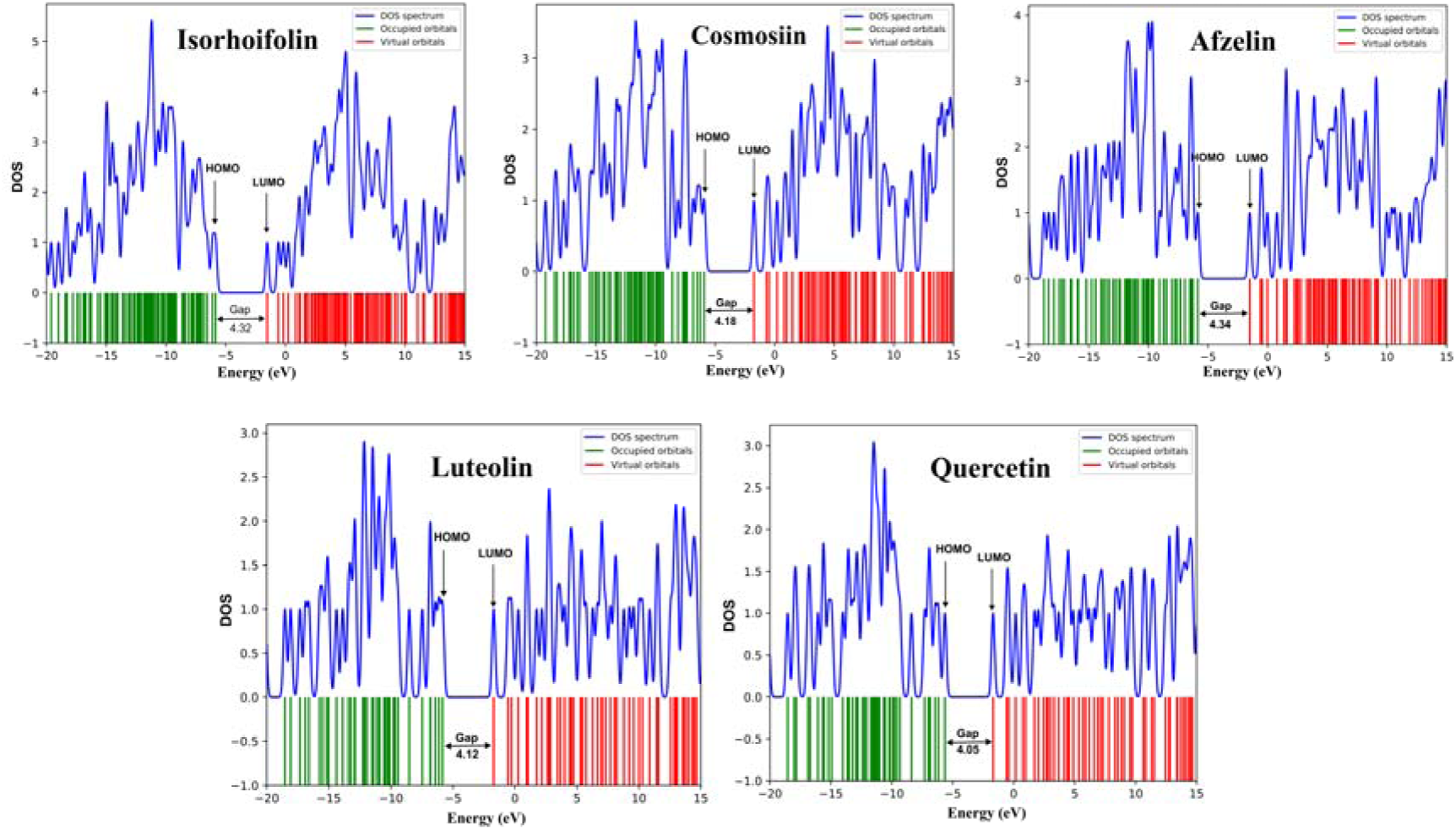
Density of state (DOS) plot of HOMO-LUMO and energy gap of five selected phytochemicals such as Isorhoifolin, Cosmosiin, Afzelin, Luteolin and Quercetin of *Cuscuta reflexa*.

### Molecular electrostatic potential analysis

The molecular electrostatic potential (MEP) analysis of the Isorhoifolin, Cosmosiin, Afzelin, Luteolin and Quercetin, provides crucial insights into the charge distribution across their surfaces. The color gradient ranges from red (indicating regions of negative electrostatic potential) to blue (representing areas of positive potential), with green and yellow zones illustrating neutral charge areas. The positive electrostatic potential varied between +6.901 a.u. and +8.347 a.u., while the negative electrostatic potential ranged from -6.901 to -8.347 a.u. Isorhoifolin shows a moderate range from -6.901 to +6.901 a.u., with negative potential around oxygen atoms (nucleophilic sites) and positive potential near hydrogen atoms (electrophilic sites). Cosmosiin exhibits a broader range, from -7.604 to +7.604 a.u., indicating stronger charge separation. Afzelin extends this range further, from -7.824 to +7.824 a.u., demonstrating greater polarity. Luteolin shows the most extreme values, from - 8.347 to +8.347 a.u., suggesting high reactivity, while Quercetin has a similar range, from - 8.232 to +8.232 a.u. The MEP range spans from -8.347 to +8.232 a.u., with Luteolin and Quercetin showing the strongest charge separation, implying high reactivity. Cosmosiin and Afzelin exhibit significant, though more balanced, reactivity, and Isorhoifolin displays moderate polarity (**Figure 9**).

**Figure 9:** The molecular electrostatic potential (MEP) maps of the selected phytochemicals like Isorhoifolin, Cosmosiin, Afzelin, Luteolin and Quercetin from *Cuscuta reflexa*.

### Vibrational frequencies analysis

Fourier Transform Infrared (FT-IR) spectroscopy is a widely used analytical technique for identifying functional groups such as hydroxyl (-OH), carboxyl (-COOH), carbonyl (C=O), alkene (C=C), and aromatic groups which can be analyzed in their infrared absorption spectra. The vibrational frequency spectrum, spanning from 400 to 4000 cm[¹, depicted in **(Table S3 and Figure 10A)**. Following, Isorhoifolin, the νC-H stretch (aromatics & alkenes) occurs at 3139 cm[¹ (scaled to 3020 cm[¹), with a notable νC=O stretch for the carbonyl group at 1717 cm[¹ (scaled to 1652 cm[¹). In Cosmosiin, similar vibrational modes are observed, with a slightly higher νC-H stretch at 3184 cm[¹ (scaled to 3063 cm[¹), and the νC=O stretch appears at 1719 cm[¹ (scaled to 1654 cm[¹). Afzelin shows a prominent νO-H stretch for alcohols at 3824 cm[¹ (scaled to 3678 cm[¹), much higher than the other compounds, alongside a νC=O stretch at 1735 cm[¹ (scaled to 1669 cm[¹), indicating a stronger carbonyl bond than in Isorhoifolin and Cosmosiin. Luteolin stands out with the highest νO-H stretch for alcohols at 3838 cm[¹ (scaled to 3692 cm[¹), indicating a very strong O-H bond, while its νC=O stretch is similar to the others at 1718 cm[¹ (scaled to 1653 cm[¹). On the other hand, Quercetin exhibits a slightly lower νO-H stretch at 3777 cm[¹ (scaled to 3633 cm[¹), along with two νC=C stretches for alkenes at 1673 cm[¹ and 1658 cm[¹ (scaled to 1609 cm[¹ and 1595 cm[¹, respectively), reflecting more complex alkene interactions. While most compounds show νC-O stretches in the 1200–1300 cm[¹ range, Afzelin and Quercetin display prominent tertiary alcohol and ester stretches, with Quercetin’s νC-O stretch for aromatic esters reaching 1299 cm[¹ (scaled to 1250 cm[¹). Notably, the νC-N stretch, characteristic of amine groups, is consistently lower, ranging from 1021 cm[¹ (Isorhoifolin) to 1332 cm[¹ (Afzelin), reflecting the weaker bond strength of these groups compared to carbonyls or hydroxyls.

**Figure 10:** Vibrational frequencies and UV-Visible spectra of identified compounds of *Cuscuta reflexa*, (A) Values of vibrational frequency (C=O, O–H, C=C, C-N, C-O, C=O and C–H stretches) are indicated by different arrows. (B) The UV-Visible spectra display the maximum absorbance (epsilon; λ_max_) values, excitation energies, and oscillator strengths of the selected compounds. Peaks of different colors correspond to specific compounds.

### UV-visible spectral analysis

To explore electronic transitions in selected five compounds, we employed TD-DFT calculations. This approach facilitated the determination of essential spectroscopic parameters such as maximum absorbance (λmax) values, excitation energies, oscillator strengths, and transition assignments (**Figure 10B**). The λmax reflects the energy of molecular orbital transitions and oscillator strengths measure transition intensity, while excitation energies measure energy requirements. Transition assignments characterize orbitals. The singlet ground state (S_0_) is the predominant state for these molecules. When they absorb light, they transition to either the singlet excited state S_1_ or S_2_ **(Table 4)**. In this study, the absorption spectrum in the experiment was taken to range between 100 to 550 nm. The molecules are depicted as their maximum absorption spectra corresponding to the electron transitions (H→L) from the excited state S_0_ to S_1_ or S_2_ such as 0.65315 (85.32 %) for Isorhoifolin, 0.68421 (93.62 %) for Cosmosiin, 0.67068 (89.96 %) for Afzelin, 0.66538 (88.54 %) for Luteolin and 0.67680 (91.61 %) for Quercetin. The peak wavelengths are observed at 329.31 nm, 345.12 nm, 346.77 nm, 348.03 nm, and 356.54 nm with excitation energies 3.7650 eV, 3.5925 eV, 3.9043 eV, 3.5625 eV and 3.4774 eV, respectively.

**Table 4:** Electronic absorption spectra of *Cuscuta reflexa* compounds calculated at TD-DFT/ B3LYP/ 6-311g (d,p)

| Name | Excited State | Assignment | Coefficient | Energy (eV) | Wavelength (nm) | Oscillator strength |
| --- | --- | --- | --- | --- | --- | --- |
| Isorhoifolin | $S_0 \rightarrow S_1$ | H-1 → L | -0.22896 (10.48 %) | 3.7650 | 329.31 | 0.1477 |
| | | H → L | 0.65315 (85.32 %) | | | |
| | $S_0 \rightarrow S_2$ | H-3 → L | 0.59612 (71.07 %) | 4.0301 | 307.64 | 0.0624 |
| | | H-1 → L | 0.33541 (22.49 %) | | | |
| Cosmosiin | $S_0 \rightarrow S_1$ | H-1 → L | 0.12206 (2.97 %) | 3.5925 | 345.12 | 0.0780 |
| | | H → L | 0.68421 (93.62 %) | | | |
| | $S_0 \rightarrow S_2$ | H-3 → L | 0.67516 (91.16 %) | 3.9562 | 313.39 | 0.0097 |
| | | H-3 → L+1 | 0.11593 (2.68 %) | | | |
| | | H-1 → L | 0.12402 (3.07 %) | | | |
| Afzelin | $S_0 \rightarrow S_1$ | H-3 → L | -0.18005 (6.48 %) | 3.5754 | 346.77 | 0.0003 |
| | | H-1 → L | 0.65062 (84.66 %) | | | |
| | | H-1 → L+1 | -0.12083 (2.91 %) | | | |
| | $S_0 \rightarrow S_2$ | H-2 → L | 0.18615 (6.93 %) | 3.9043 | 317.55 | 0.2804 |
| | | H → L | 0.67068 (89.96 %) | | | |
| Luteolin | $S_0 \rightarrow S_1$ | H-1 → L | -0.21620 (9.34 %) | 3.5625 | 348.03 | 0.1261 |
| | | H → L | 0.66538 (88.54 %) | | | |
| | $S_0 \rightarrow S_2$ | H-4 → L | -0.46143 (42.58 %) | 3.9445 | 314.32 | 0.1401 |
| | | H-1 → L | 0.48927 (47.87 %) | | | |
| | | H → L | 0.14672 (4.30 %) | | | |
| Quercetin | $S_0 \rightarrow S_1$ | H-1 → L | 0.16881 (5.69 %) | 3.4774 | 356.54 | 0.2288 |
| | | H → L | 0.67680 (91.61 %) | | | |
| | $S_0 \rightarrow S_2$ | H-2 → L | 0.12896 (3.32 %) | 4.0082 | 309.32 | 0.2483 |
| | | H-1 → L | 0.65982 (87.07 %) | | | |
| | | H → L | -0.15721 (4.94 %) | | | |

## Discussion

Sequence analysis of LVDD9B offers hypothetical insights into its role as a dual cytoplasmic and possibly extracellular cuticular protein. The tools such as CELLO strongly elucidate this cytoplasmic subcellular localization of LVDD9B cuticular protein since it will have a central role in protein synthesis-cum-cellular signaling [43]. However, the extracellular prediction by ProtParam proposes dual functionality, which was commonly observed in similar proteins from crustaceans whereby their extracellular presence was related to structural assembly while cytoplasmic functions were cellular stress responses [44]. Thus, such evolutionary advantages could be achieved by the ability of the protein to perform many biological functions during cuticle formation, repair, and immune reactions. The critical structural integrity feature of LVDD9B is evidenced by the conserved chitin-binding site. The chitin-binding domains anchor proteins within the chitin matrix, which strengthens the exoskeleton against environmental stress and pathogenic invasion [45]. Predictions about secondary structures have demonstrated how LVDD9B is based on beta strands and an alpha helix to maintain structural integrity and flexibility. Beta strands have been associated with chitin-binding proteins which further harbor polysaccharide interaction with cuticles. An alpha helix maintains the protein in a stable position since this has been observed with studies that consider proteins under environmental stress [46]. Preparation of three-dimensional structure predictions using SWISS-MODEL, validated effectively through many high-confidence approaches like ProSA Z-scores or Ramachandran plots, covers the more comprehensive interpretation of structural integrity on LVDD9B. Ninety-one percent of residues in the highly favored areas would fit well in an alignment with other well-characterized chitin-binding proteins shows in the Ramachandran plot at the Procheck algorithm, thus supporting this consideration from the functional standpoint as well [47]. Docks of LVDD9B with viral protein VP26 just opened the doors for understanding what molecular mechanisms would be used in shrimp-virus interactions and assured critical clues to pathogen exploitation of host proteins. The docking study had the HADDOCK score of -99.9 ± 9.5, and thus sites a very strong interface for interaction. That huge contribution of electrostatic (-263.1 ± 34.8 kcal/mol) and van der Waals energies (-79.7 ± 9.3 kcal/mol) stresses further the role of non-covalent interactions in stabilization of the protein complex. The buried surface area (BSA) of 2363.1 Å² additionally complements the functional import of the LVDD9B-VP26 interaction. Large BSAs are indicative of a high degree of binding affinity in functional protein complexes[48], confirming their contribution to the interaction through Arg28, Asp72, Arg103, Lys79, Asn80, and Ala180. Residuals thus act as possible hot spots for any therapeutic intervention of the targeted these binding sites on the viral replication inhibitors. These findings suggest that this interaction could serve as a LVDD9B potential target for future inhibitors against WSSV in shrimp. Plant-derived bioactive compounds are gaining traction as viable therapeutic alternatives for WSSV management. *Cuscuta reflexa* offers a spectrum of pharmacological activities, including antispasmodic, antiviral, antibacterial, antifungal, and antitumor properties, among others [49]. This study targets the identification of bioactive molecules from *Cuscuta reflexa* capable of inhibiting LVDD9B. These compounds were subjected to comprehensive computational evaluations, encompassing thermodynamic stability, molecular docking, interaction patterns, ADMET profiling, MDS, and post-MDS validations to determine their efficacy against LVDD9B. Conceiving pharmacokinetics of luteolin, quercetin, cosmosiin, isorhoifolin give manifold conclusions about their therapeutic usage and environmental safety, bearing out the practical implementation in aquaculture. Of all these compounds, Luteolin, and Quercetin exhibit preferred pharmacokinetics according to Lipinski’s Rule of Five (L5) criteria. All of the investigated compounds demonstrate a 30 – 70% absorption rate that reflects moderate absorption, while luteolin indicates 81% absorption rate[50]. However, selected phytocompounds did not inhibit the function of any of the several metabolic substrates or inhibitors in hepatocytes indicating proper metabolism. Importantly, the renal excretion profile was favorable, with no indication of toxic buildup due to negative OCT2 interactions [51]. On top of that, the LD50 reading for oral rat acute toxicity of these substances fell within an acceptable range of 2 to 2.5, presenting that compounds are harmless for human ingestion [52]. The ecological risk assessment of the selected phytochemicals indicates great differences in their impact on the environment, pointing out the need for an approach that balances therapeutic efficacy with ecological safety [53]. From all the compounds investigated, Cosmosiin and Isorhoifolin are the most environmentally friendly one than other phytochemicals. Additionally, aquatic organisms like fish toxicity range of 0.34 to 0.53 and crustacean toxicity range 0.12 to 0.25, suggesting no toxicity of selected phytocompounds. However, Cosmosiin, Afzelin, Luteolin and Quercetin exhibits highly toxicity to *P. subcapitata* and *T. pyriformis*. For terrestrial organism such as honeybees, quail (*Colinus virginianus*), and duck (*Anas platyrhynchos*) all of the studied ligands exhibit no toxicity, indicate All of compounds are holds out an empty-handed ecological impact for terrestrial ecosystems [54]. Higher biodegradability rates fast breakdown the bioactive compounds and engage long term ecological degradation. Conversely, lower biodegradability gives a worrying aspect of environmental persistence that could bring about long exposure risks in both aquatic and terrestrial settings [55]. Among the tested compounds, Cosmosiin and Luteolin exhibit higher biodegradability, whereas Quercetin demonstrates lower biodegradability. In our docking study we accepted the top five compounds plus a control drug with the highest binding affinities observed were -8.5, -8, -7.3, -7.2 and 7.1 Kcal/mol for Isorhoifolin, Cosmosiin, Afzelin, Luteolin, and Quercetin, respectively. Based on these findings, we conducted further molecular interaction analysis to determine how molecules interact with favorable interacting sites of protein residues. All of the compounds create strong connections with the LVDD9B protein through the formation of five hydrogen bonds, as depicted in **(Figure 3** and **Table S1).** MDS provide a real-time window into the conformational shifts that proteins undergo upon ligand binding. In our study, trajectory analyses revealed subtle conformational adjustments in the protein structure upon interaction with the ligands. These changes were minimal, signifying that the ligands stabilize the protein’s structure, which is critical for maintaining its functional integrity. Such conformational insights offer a better understanding of how ligands may affect protein behavior in a dynamic biological environment [37]. The RMSD value helps assess the average positional changes of selected atoms across a given reference timeframe. In our study exhibited that ligands with minimal RMSD fluctuations were observed, where the hypothetical nature of the protein structure caused the RMSD values to slightly exceed the acceptable range (1-3 Å). The visualization of the trajectory showed that the bioactive compounds stayed attached to the binding site of the receptor protein throughout the entire simulation. This consistent interaction suggests a strong affinity between compounds and the protein, potentially enhancing their therapeutic efficacy [56]. RMSF data also verifies that the complex’s low fluctuation designates the compounds’ stability within the targeted receptor, whereas higher fluctuations indicate less stability within the protein [57]. Our findings showed that the RMSF data confirm that the stability of the compounds with the targeted receptor is determined by the complex’s minimal fluctuation. The center of mass from the protein C terminals and N terminals examines the stability of the protein structure. It gives a vast assessment of protein folding characteristics for interpreting the Rg value. A higher Rg value represents the dissociation of the compounds from the receptor, while a lower Rg value indicates an increase in compactness [58]. In this study, the selected compounds provided notable compactness with the protein. The lowest SASA values observed in our study indicate strong hydrophilic interactions between the ligands and the surrounding solvent [59]. These interactions are vital for maintaining the stability and functionality of the protein-ligand complex in biological systems. Similarly, MolSa and PSA were analyzed and the outcomes efficiently corroborate the data. Ligands are engaged in a variety of with the protein receptor during MDS such as hydrophobic interactions, hydrogen bonds, metals, ionic bonds and water bridge. The stability of these interactions throughout the simulation was key to establishing a firm protein-ligand binding and ensuring the reliability of the molecular interactions. Furthermore, we employed the MM-GBSA calculation to determine the binding free energy. The higher ΔG value represent more tighter interaction with the target protein (**Figure 5**), while the investigated compounds are showing effective results [60]. In the PCA study, each dot represents an individual frame of the simulation. The PCA scatter plot revealed that the collective motions of the protein – ligand complexes did not deviate significantly from the apo-protein, occupying less conformational space that indicates stable binding of the ligands. This stability indicates the compounds may induce a favorable environment for protein, potentially enhancing its functional efficiency [61]. To gain a deeper understanding of the thermal efficiency of the investigated compounds, we conducted density functional theory calculations using Gaussian. The studied phytocompounds are comparatively exhibited lower hardness and higher softness. In addition, it was indicated that the experimental compounds exhibited greater stability and had potential interactions with the target receptor [62]. Among the ligands, Quercetin was found to have the lowest hardness value (2.02 eV) and the highest softness value (0.49 eV), indicating that it is more reactive than other investigated compounds [63]. Furthermore, the HOMO-LUMO gap obtained from frontier molecular analysis helps determine a compounds stability and its reactivity with the receptors [64]. Afzelin demonstrated a higher energy gap, suggest increased stability and reduced reactivity, while Quercetin exhibited lower energy gap, indicates heightened reactivity. This contrast stems from their electron-donor or electron-acceptor properties, where a smaller energy gap facilitates more effective interactions with receptors [65]. The order of energy gaps can be expressed as Afzelin < Isorhoifolin < Cosmosiin < Luteolin < Quercetin. In Molecular electrostatic potential (MEP) analysis, Quercetin and Luteolin showed positive potentiality toward oxygen, while Cosmosiin and Afzelin exhibit significantly moderate reactivity toward oxygen and hydrogen atoms. Among them Isorhoifolin showed highest electrostatic potential for nucleophilic sites and lowest potential for electrophilic sites [66]. However, Vibrational frequencies also calculated of the studied compounds to determine the functional groups and closely compare with the experimental FT-IR values for confirmation of the presence of functional groups within the structures. Nevertheless, aromatics carbonyls, alkene, aromatics, esters, amines alcohol and so on functional groups are identified through the FT-IR spectrum analysis [67]. In studying the UV spectra, various features such as excitation energy, oscillator strength and absorption wavelengths of selected ligands were examined. The absorption spectrum in this experiment ranged from 100 to 550 nm and reveals a fascinating dance of electrons. These electrons jump from the highest occupied molecular orbital (HOMO) to the lowest unoccupied molecular orbital (LUMO) [68]. The highest absorption in the gaseous phase of the compounds occurred at 329 nm (3.76 eV), 345 nm (3.6 eV), 347 nm (3.6 eV), 348 nm (3.56 eV) and 356 nm (3.5 eV) due to H → L (85 %), H → L (93 %), H-1 → L (85 %), H → L (88.5%) and H → L (92 %) transitions, while experimental absorption of UV-Visible was recorded in Cosmosiin, Afzelin, Luteolin, and Quercetin and it highest absorption peaks were observed at 330 nm, 360 nm, 330 nm and 374 nm, respectively [69–71]. This small variation in wavelength absorption across different compounds indicates that these ligands exhibited high binding affinity to their target receptors. Computational evaluations of the distinct properties of the five compounds, supported by a range of results, guided the selection of these compounds for further assessment via in vivo studies and in vitro cell line investigation.

## Conclusion

In conclusion, this study highlights the structural and functional properties of the LVDD9B protein, mentioning its potential role in immune responses and cuticle formation in shrimp. Molecular docking and dynamics simulations revealed strong interactions between LVDD9B and various bioactive compounds from *Cuscuta reflexa* plant, particularly Luteolin and Quercetin, which exhibited favorable pharmacokinetic profiles. The ecological risk assessment showed Cosmosiin as the least hazardous compound, with low toxicity and high biodegradability, suggesting its suitability for sustainable aquaculture practices. However, Quercetin’s potential for bioaccumulation in aquatic ecosystems raises concerns for its environmental impact. Future research should focus on optimizing Cosmosiin’s therapeutic potential, enhancing its efficacy, and evaluating its long-term safety. Further in vitro and in vivo studies are necessary to confirm the bioactivity of these compounds and explore their practical applications in antiviral therapies for shrimp diseases like WSSV. Additionally, improving the environmental safety of other compounds, such as Quercetin, is essential for their broader use in aquaculture.

## Authors Contribution

**Md. Iftehimul:** Conceptualization, Methodology, Data curation, Formal analysis, Writing - original draft, Writing - review & editing. **Neaz A. Hasan:** Conceptualization, Writing - original draft, Writing - review & editing. **Mst. Farzana Akter:** Writing – original draft, Methodology. **Md. Arju Hossain** and **Sajia Afrin Tima:** Writing – original draft, review and editing**. Amirul Kabir:** Writing – original draft, Methodology. **Prottay Choudhury:** Formal analysis, Visualization. **Apurbo Bhowmick:** Methodology, Visualization. **Sakib Anzum Pranto:** Methodology, Data curation, Visualization. **Ali Mohamod Wasaf Hasan** and **Siddique Akber Ansari:** Writing - review & editing. **Md Habibur Rahman:** Conceptualization, Methodology, and Supervision.

