## Supplement File for "In Silico Evaluation and Therapeutic Targeting of LVDD9B Protein for WSSV Inhibition: Molecular and Ecological Insights for Aquaculture Solutions"

**Figure S1:** Prediction of Secondary structure of LVDD9B protein through (A) PSIPRED and (B) JPred.

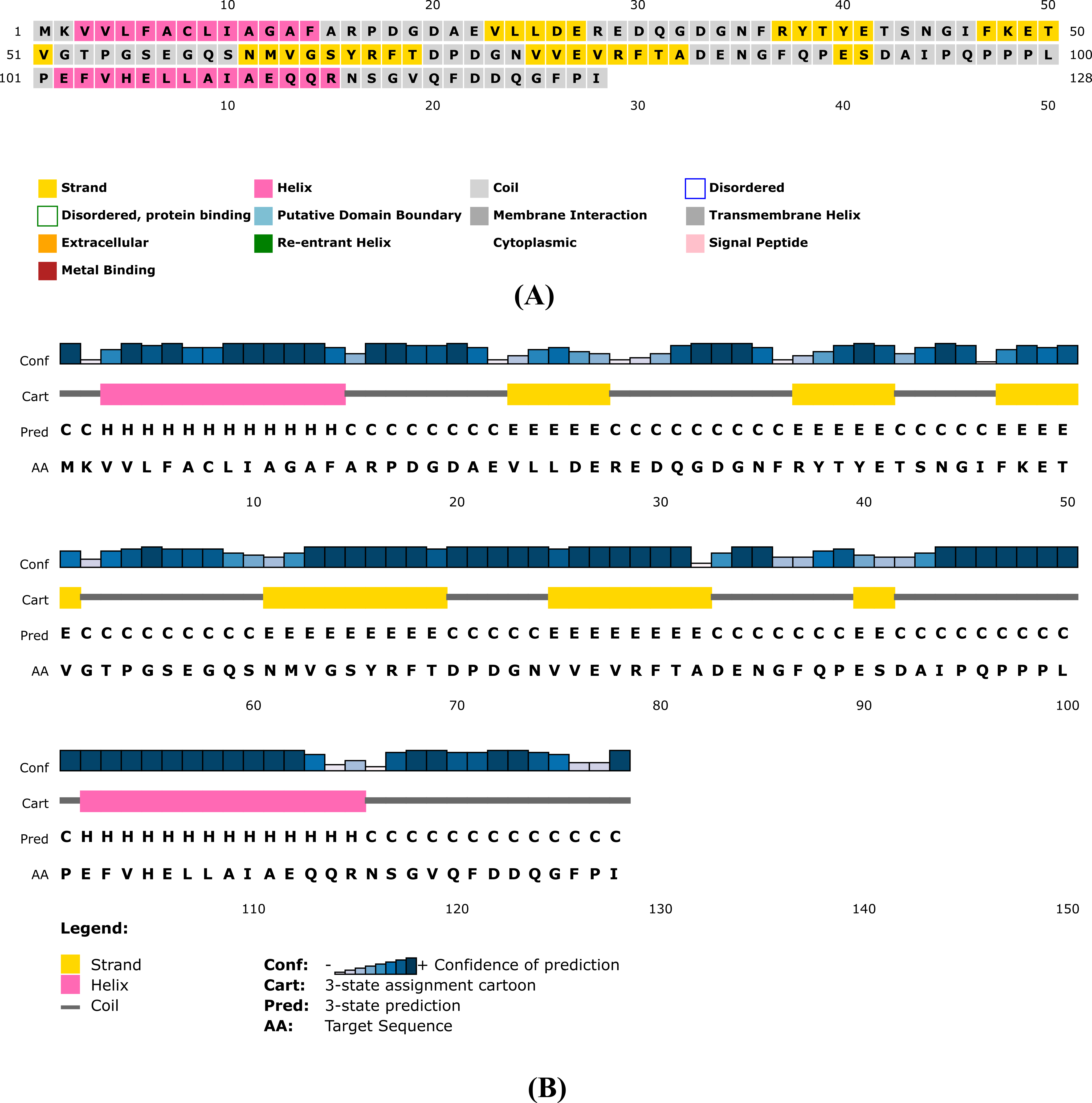

**Figure S2:** Validation of Tertiary structure of LVDD9B protein using (A) Prosa- web, (B) Verify3D and (C) Procheck.

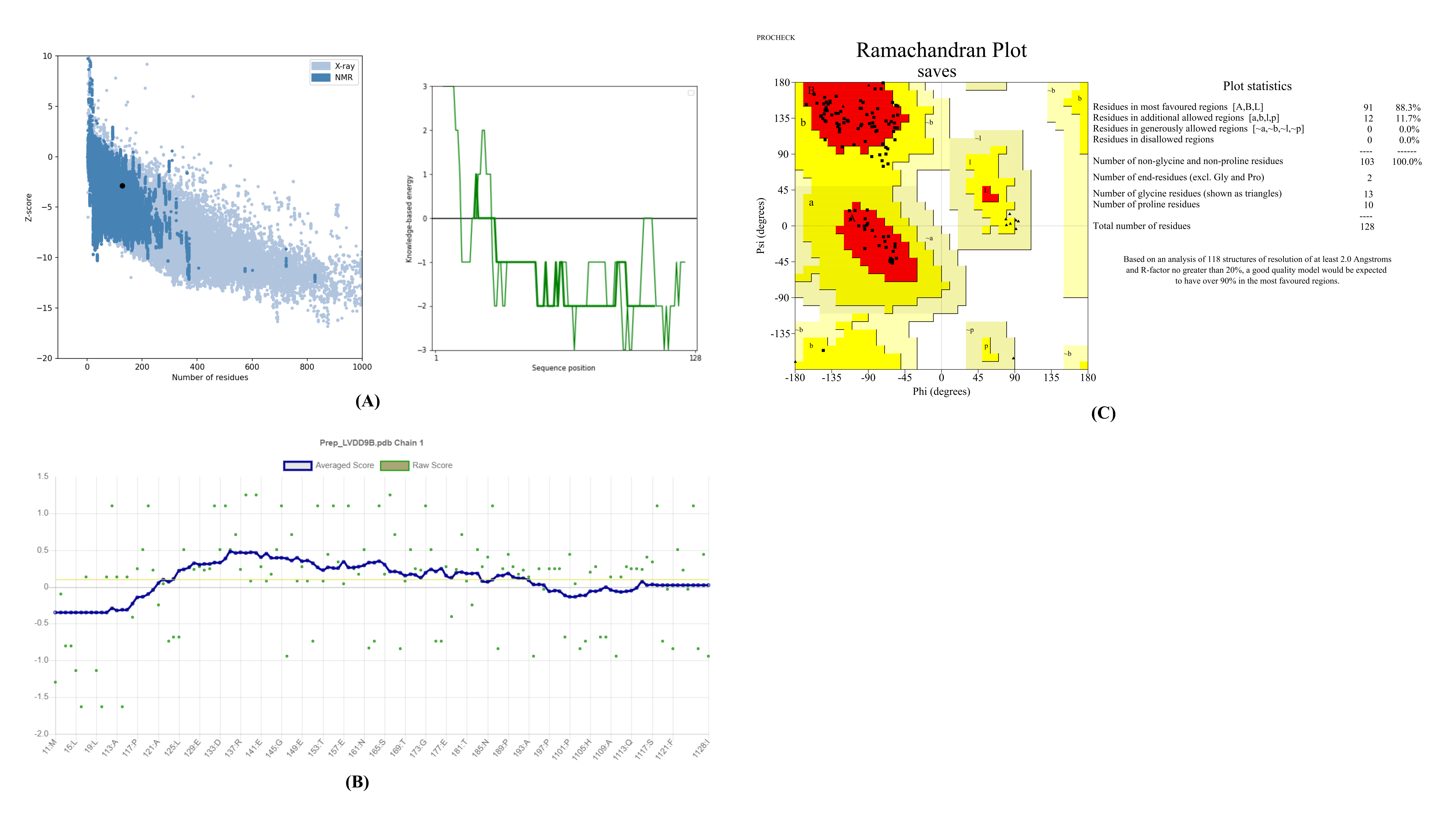

**Figure S3:** Post simulation Intermolecular interactions of LVDD9B and phytocompounds.

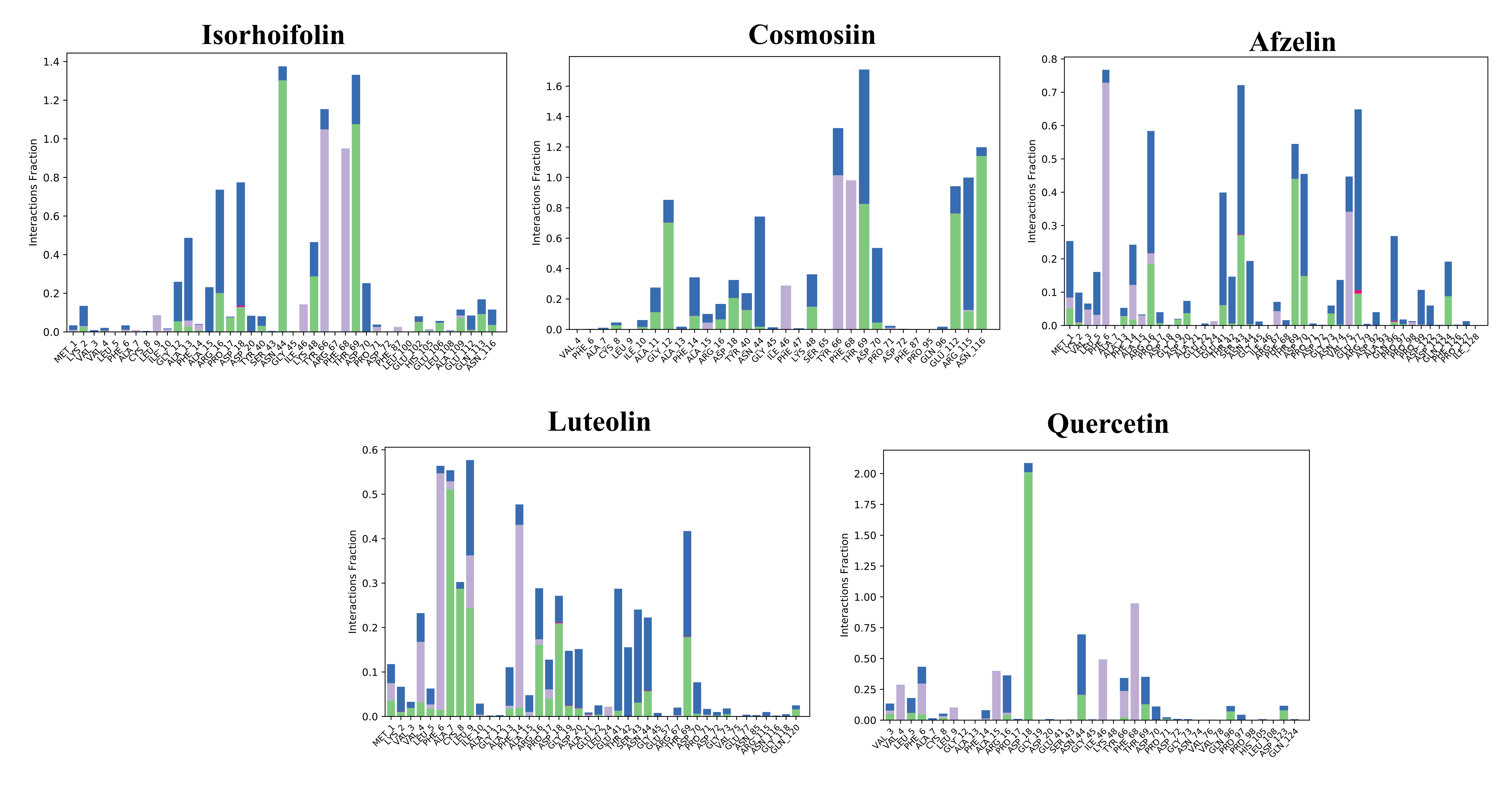

**Figure S4:** HOMO-LUMO distribution plots of *Cuscuta reflexa* phytocompounds in the ground state at DFT in the gas phase.

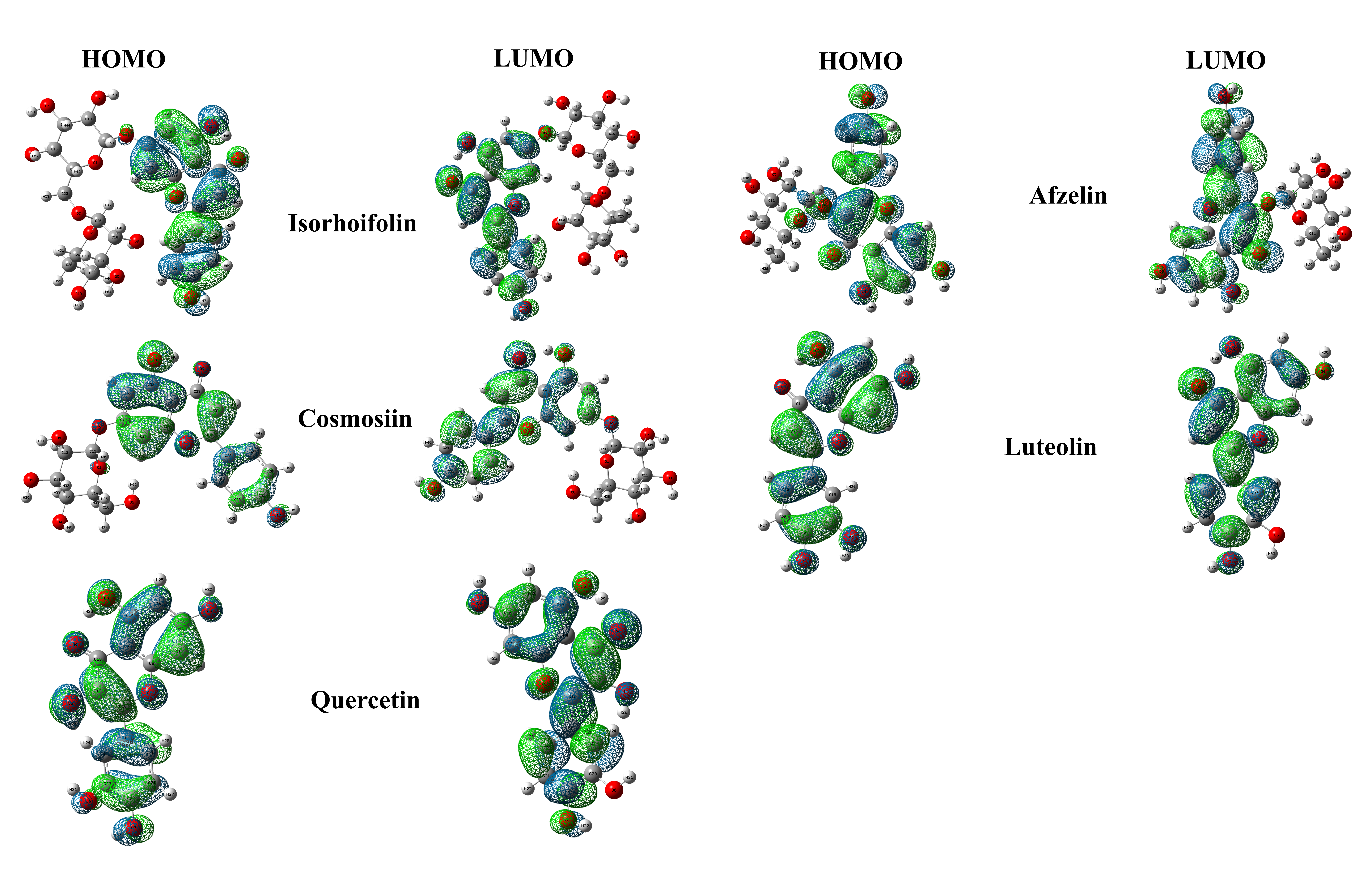

**Table S1:** Interpretation of Molecular Interaction of phytocompounds and Targeted Receptor.

| Protein – Ligand Interaction | Binding energy (kcal/mol) | H-bond residues | H-bond length (Å) | No. of H-bonds | Other bond residues |
| --- | --- | --- | --- | --- | --- |
| Isorhoifolin_LVDD9B | -8.5 | Phe14 | 3.00 | 5 | Ala13, Asp72, Gly73, Asp70, Thr69, Asp18, Tyr66, Ile46, Phe68 |
|  |  | Pro71 | 3.21 |  |  |
|  |  | Tyr40 | 2.85 |  |  |
|  |  | Asn44 | 2.84 |  |  |
|  |  |  | 2.87 |  |  |
| Cosmosiin_LVDD9B | -8 | Asp70 | 2.93 | 5 | Tyr66, Ile46, Phe68, Asp18, Thr69 |
|  |  | Pro71 | 3.20 |  |  |
|  |  | Tyr40 | 2.76 |  |  |
|  |  | Asn44 | 3.01 |  |  |
|  |  |  | 2.92 |  |  |
| Afzelin_LVDD9B | -7.3 | Asp70 | 3.14 | 5 | Pro71, Arg16, Asp18, Ser43, Gly73, Gly45 |
|  |  | Pro17 | 3.24 |  |  |
|  |  | Asp20 | 3.25 |  |  |
|  |  | Asn44 | 2.86 |  |  |
|  |  | Thr69 | 3.01 |  |  |
| Luteolin_LVDD9B | -7.2 | Asp20 | 3.17 | 5 | Pro71, Arg16, Asn44, Ser43, Asp18 |
|  |  | Pro17 | 3.14 |  |  |
|  |  | Thr69 | 2.95 |  |  |
|  |  | Asp70 | 3.06 |  |  |
|  |  |  | 2.80 |  |  |
| Quercetin_LVDD9B | -7.1 | Thr69 | 2.93 | 5 | Pro71, Asp18, Arg16, Ser43, Asn44 |
|  |  | Asp20 | 3.19 |  |  |
|  |  | Pro17 | 3.14 |  |  |
|  |  | Asp70 | 3.07 |  |  |
|  |  |  | 2.81 |  |  |

**Table S2:** Pharmacokinetics properties determination of the selected Hits

| Properties | Action | Isorhoifolin | Cosmosiin | Afzelin | Luteolin | Quercetin |
| --- | --- | --- | --- | --- | --- | --- |
| Physiochemical  Properties | Molecular Weight (g/mol) | 578.52 | 432.38 | 432.38 | 286.24 | 302.24 |
|  | Number of rotatable bonds | 6 | 4 | 3 | 1 | 1 |
|  | Hydrogen bond acceptors | 14 | 10 | 10 | 6 | 7 |
|  | Hydrogen bond donors | 8 | 6 | 6 | 4 | 5 |
|  | Molar Refractivity | 137.33 | 106.11 | 106.97 | 76.01 | 78.03 |
|  | TPSA (A^2^) | 228.97 | 170.05 | 170.05 | 111.13 | 131.36 |
|  | Consensus Log P (o/w) | -0.41 | 0.55 | 0.60 | 1.73 | 1.23 |
|  | Lipinski violation | Yes (3) | Yes (1) | Yes (1) | no | no |
|  | Bioavailability score | 0.17 | 0.55 | 0.55 | 0.55 | 0.55 |
| Absorption | Water solubility (log mol/L) | -2.942 | -2.559 | -2.969 | -3.094 | -2.925 |
|  | CaCo2 permeability  (log Papp in 10^-6^ cm/s) | 0.285 | 0.33 | 0.123 | 0.096 | -0.229 |
|  | Intestinal absorption (human) (% absorbed) | 32.393 | 37.609 | 60.006 | 81.13 | 77.207 |
|  | Skin permeability (log Kp) | -2.735 | -2.735 | -2.735 | -2.735 | -2.735 |
|  | P-Glycoprotein I and II inhibitor | no | no | no | no | no |
| Distribution | VDss (Human) (Log L/kg) | 1.815 | 0.342 | 1.15 | 1.153 | 1.559 |
|  | Fraction unbound (Human) (Fu) | 0.116 | 0.218 | 0.087 | 0.168 | 0.206 |
|  | BBB permeability (log BB) | -1.72 | -1.391 | -1.265 | -0.907 | -1.098 |
|  | CNS permeability (log PS) | -4.727 | -3.746 | -3.994 | -2.251 | -3.065 |
| Metabolism | CYP2D6 Substrate | no | no | no | no | no |
|  | CYP3A4 Substrate | no | no | no | no | no |
|  | CYP2C9 Inhibitor | no | no | no | no | no |
|  | CYP2C19 Inhibitor | no | no | no | no | no |
|  | CYP2D6 Inhibitor | no | no | no | no | no |
|  | CYP3A4 Inhibitor | no | no | no | no | no |
| Excretion | Total clearance (log/ml/min/kg) | -0.016 | 0.547 | 0.431 | 0.495 | 0.407 |
|  | Renal OCT2 substrate | no | no | no | no | no |
| Toxicity | AMES toxicity | no | no | no | no | no |
|  | Max. Tolerated Dose (Hum) (log mg/kg/day) | 0.533 | 0.515 | 0.544 | 0.499 | 0.499 |
|  | hERG I inhibitor | no | no | no | no | no |
|  | hERG I I inhibitor | yes | no | yes | no | no |
|  | Skin Sensitization | no | no | no | no | no |
|  | Oral Acute Toxicity/LD50 (mol/kg) | 2.52 | 2.595 | 2.61 | 2.455 | 2.471 |
|  | Oral chronic toxicity (log mol/ kg/day) | 3.338 | 4.359 | 2.799 | 2.409 | 2.612 |
|  | Hepatotoxicity | no | no | no | no | no |

**Table S3:** Selected vibrational frequencies (cm^-1^) of *Cuscuta reflexa* compounds were calculated in the gas phase (scaled).

| Name | Assignments | Functional Group | Vibrational Frequencies  (cm-1)  (Initial) | Vibrational Frequencies  (cm-1)  (Scaled) |
| --- | --- | --- | --- | --- |
| Isorhoifolin | *ν*C-H stretch | Aromatics & alkenes | 3139 | 3020 |
|  | νC=O stretch | Carbonyls | 1717 | 1652 |
|  | *ν*C=C stretch | Alkenes | 1659 | 1596 |
|  | *ν*C-C stretch | Aromatics (in–ring) | 1542 | 1483 |
|  | *ν*C-O stretch | Tertiary alcohol | 1208 | 1162 |
|  | *ν*C-N stretch | Amines | 1061 | 1021 |
| Cosmosiin | *ν*C-H stretch | Aromatics | 3184 | 3063 |
|  | νC=O stretch | Carbonyls | 1719 | 1654 |
|  | *ν*C=C stretch | Alkene | 1662 | 1599 |
|  | *ν*C-C stretch | Aromatics (in–ring) | 1543 | 1484 |
|  | *ν*C-N stretch | Aromatic amine | 1377 | 1325 |
|  | *ν*C-O stretch | Esters | 1210 | 1164 |
|  | *ν*C-N stretch | Amines | 1083 | 1042 |
| Afzelin | *ν*O-H stretch | Alcohol | 3824 | 3678 |
|  | *ν*C-H stretch | Aromatics & alkenes | 3047 | 2931 |
|  | νC=O stretch | Carbonyls | 1735 | 1669 |
|  | *ν*C=C stretch | Alkenes | 1662 | 1599 |
|  | *ν*C-C stretch | Aromatics (in–ring) | 1494 | 1437 |
|  | *ν*C-N stretch | Aromatic amine | 1385 | 1332 |
|  | *ν*C-O stretch | Esters | 1234 | 1187 |
|  | *ν*C-O stretch | Tertiary alcohol | 1181 | 1136 |
| Luteolin | *ν*O-H stretch | Alcohol | 3838 | 3692 |
|  | *ν*C-H stretch | Aromatics & alkenes | 3142 | 3023 |
|  | νC=O stretch | Carbonyls | 1718 | 1653 |
|  | *ν*C=C stretch | Alkenes | 1669 | 1606 |
|  | *ν*C-C stretch | Aromatics (in–ring) | 1556 | 1497 |
|  | *ν*C-N stretch | Aromatic amine | 1346 | 1295 |
|  | *ν*C-O stretch | Tertiary alcohol | 1179 | 1134 |
|  | *ν*C-O stretch | aliphatic ether | 1143 | 1100 |
| Quercetin | *ν*O-H stretch | Alcohol | 3777 | 3633 |
|  | *ν*C-H stretch | Aromatics & alkenes | 3171 | 3051 |
|  | νC=O stretch | Aldehyde | 1714 | 1649 |
|  | *ν*C=C stretch | Alkenes | 1673 | 1609 |
|  | *ν*C=C stretch | Alkenes | 1658 | 1595 |
|  | *ν*C-C stretch | Aromatics (in–ring) | 1556 | 1497 |
|  | *ν*C-N stretch | Aromatic amine | 1340 | 1289 |
|  | *ν*C-O stretch | Aromatic ester | 1299 | 1250 |
|  | *ν*C-O stretch | Esters | 1226 | 1179 |
|  | *ν*C-O stretch | Tertiary alcohol | 1176 | 1131 |
